# Mechanistic basis of EMRE’s essential role in the regulation of mitochondrial calcium uniporter complex

**DOI:** 10.64898/2026.06.25.733848

**Authors:** Anshu Kumari, Dung M. Nguyen, Deborah Disilvestre, Nathaniel D. A. Dirda, Sri Harsha Kethanapalli, Joseph P. Y. Kao, Vivek Garg

## Abstract

Mitochondrial Ca^2+^ uptake through the mitochondrial calcium uniporter complex (MCUcx) is a critical determinant of cellular metabolism, integrating Ca^2+^ signaling with ATP production and redox control. Yet how MCUcx activity is constrained to prevent Ca^2+^ overload and cell injury, and how the essential MCU regulator (EMRE), a subunit required for channel activity, mechanistically supports MCUcx function remains incompletely defined. Here, using a newly developed high-sensitivity assay to quantify MCUcx function in intact mitochondria, we uncover two fundamental roles of EMRE. First, EMRE is required for robust matrix Ca^2+^-dependent inhibition of MCUcx, acting through a juxtamembrane site via a mechanism distinct from MICU1-mediated inhibition at low cytosolic Ca^2+^. Second, by decoupling channel function from regulation, we demonstrate that EMRE promotes robust ion permeation through MCUcx, elevating its role from a structural scaffold to an active determinant of channel throughput. Together, our findings refine current models of mitochondrial Ca^2+^ regulation, establish EMRE as an essential multifunctional regulator of uniporter activity, and highlight the utility of our assay for probing MCUcx biophysical mechanisms and enabling the discovery of uniporter modulators.

**Significance Statement:** Mitochondria use Ca^2+^ signals to adjust energy production to cellular demand, but excessive Ca^2+^ entry can trigger cell death. How the mitochondrial calcium uniporter balances these opposing needs remains fundamentally unresolved. Using a high-sensitivity approach that isolates uniporter permeation from Ca^2+^-dependent confounders in intact mitochondria, we characterize a matrix Ca^2+^-dependent inhibitory mechanism that depends on EMRE and is functionally distinct from MICU1-mediated regulation. We further show that EMRE, a small regulatory subunit unique to higher organisms, not only enables channel function but promotes robust ion permeation through the pore. Together, these findings refine current models of mitochondrial Ca^2+^ regulation and provide a unified framework for understanding EMRE-dependent uniporter regulation in intact mitochondria.

## INTRODUCTION

MCUcx is embedded in the inner mitochondrial membrane (IMM) where it forms the principal pathway for Ca^2+^ transport from the cytoplasm to the mitochondrial matrix. MCUcx-mediated Ca^2+^ uptake increases metabolic output by enhancing the rate of ATP production in response to the energy demands of skeletal muscle, heart, and neurons (1–5). This is achieved by allosterically potentiating the activity of key enzymes in the Krebs cycle, particularly the pyruvate dehydrogenase complex (3, 6–8). However, excessive mitochondrial Ca^2+^ uptake via MCUcx, such as during cardiac ischemia–reperfusion injury, can lead to mitochondrial permeability transition (a pathological increase in IMM permeability) and subsequently cell death(9–12). Indeed, patients with mutations in MCUcx proteins have severe neurological, developmental and myopathic disabilities (13–16).

The exquisite selectivity of MCUcx for Ca^2+^ (*K*_d_ < 2 nM), and its ability to detect the small (∼400–700 nM), transient (0.1–5 s) elevations in cytosolic [Ca^2+^] that occur during signaling events, enable the massive and rapid mitochondrial Ca^2+^ influx that is essential for optimal metabolic control and cell viability (17, 18). The central pore of MCUcx, comprising a tetrameric assembly of MCU subunits, is tightly regulated by a number of different regulatory proteins and mechanisms. In higher eukaryotes, the pore requires an additional EMRE subunit. This single-pass membrane protein binds to each MCU subunit in a 1:1 stoichiometry and encircles the periphery of the MCU pore to create a functional channel. EMRE additionally facilitates the tethering of MICU regulatory subunits (19–21), which are also associated with the mitochondrial contact site and cristae organizing system (MICOS). MICU1-3 are EF-hand domain-containing proteins that homo- or heterodimerize and bind to MCU subunits via the intermembrane space (IMS)-exposed C-terminus of EMRE, where they modulate channel gating (along with their recently discovered binding partner, EFHD1(22)). Structures of MCUcx have revealed that MICU1 loosely associates with the pore entrance (23, 24), where it is ideally placed to impede Ca^2+^ uptake at resting cytosolic [Ca^2+^] and permit uptake when its EF-hands are occupied by Ca^2+^ (22, 23, 25, 26). However, interpreting the enhanced Ca²⁺ uptake observed in MICU1-KO cells at low cytosolic [Ca^2+^] is complicated by emerging evidence that MICU1 also has MCU-independent functions, including interactions with cristae-associated proteins and regulation of cristae morphology (17, 27–29). Together, the accessory subunits exert tight temporal and spatial control of ion flow through MCUcx in response to cellular signaling events.

Despite the significant mechanistic insights that have been gained from elegant structural and functional studies of MCUcx (17–19, 21, 23, 25, 30, 31), many aspects of regulation remain unresolved. For example, because the closed state structure of MCUcx has yet to be solved, the mechanisms underlying channel gating are not fully understood. Indeed, although the role of MICU1 as a Ca^2+^-dependent gatekeeper of MCUcx is well-supported, the contributions of other Ca^2+^-dependent binding partners and Ca^2+^-sensitive regulatory mechanisms require further elucidation. Previous studies have also reported matrix Ca^2+^-dependent regulation of MCU activity (32–34), although the underlying mechanism remains debated. Furthermore, the precise role of EMRE in facilitating MCU-mediated ion flux and its evolutionary advantage in metazoans remain unclear, particularly as MCU can function independently of EMRE in more primitive eukaryotes (35, 36).

A major obstacle to resolving these questions is the lack of tools to disentangle direct effects on Ca^2+^ flux through MCUcx from the broader context of Ca^2+^-dependent mitochondrial processes (30, 37–40). Mitochondria possess an intricate system to maintain matrix free [Ca^2+^] and the extramitochondrial Ca^2+^ setpoint (the cytosolic [Ca^2+^] beyond which mitochondria actively accumulate Ca^2+^) (41–43). In addition to MCUcx-dependent and -independent Ca^2+^ influx (44, 45), there are highly efficient Ca^2+^ efflux mechanisms, including a Na^+^/Ca^2+^ exchanger and a H^+^/Ca^2+^ exchanger (40), along with the high-capacity Ca^2+^ buffering provided by matrix phosphate. Measurements of MCUcx activity that use Ca^2+^ as the permeating ion must account for the effect of all these factors on matrix free [Ca^2+^] and therefore the electrochemical gradient for Ca^2+^. Mitoplast patch-clamp recordings are the most reliable method to isolate and quantitatively characterize mitochondrial ion channels in their native environment as they permit precise control over pH, ion, and voltage gradients across the IMM. Despite these advantages, however, mitoplast patch clamp involves destruction of the outer mitochondrial membrane (OMM), disruption of IMM cristae, and loss of the matrix and IMS milieu (46).

Here we develop a robust and versatile assay that utilizes Na^+^ as the permeant ion to isolate MCUcx-mediated ion uptake and its regulatory mechanisms from other Ca^2+^-dependent processes in fully intact mitochondria. Because the approach enables quantification of MCUcx function with high sensitivity and temporal resolution, and can be readily combined with other imaging and genetic tools, we have been able to uncover fundamental insights into MCUcx molecular physiology. Using this strategy, we identify a matrix Ca^2+^-dependent inhibitory mechanism of MCUcx that is mechanistically distinct from previously described modes of MCUcx regulation. We further demonstrate that this inhibitory pathway critically depends on EMRE and a conserved juxtamembrane loop within MCU. Finally, we show that EMRE directly enhances ion permeation through the MCU pore, consistent with an essential role in stabilizing the open state of the channel. Together, these findings redefine EMRE as a multifunctional regulator of mitochondrial Ca^2+^ uptake and establish a broadly applicable experimental platform for probing MCUcx biophysics, and the functional consequences of disease-linked variants and pharmacological modulators.

## RESULTS

### Ionomycin enables *in situ* measurements of MCUcx activity

Because Ca^2+^ is both the primary permeating ion and a regulator of MCUcx, we chose Na^+^ as the permeating ion to study regulation of MCUcx without the confounding influences of permeating Ca^2+^. Na^+^ has the same ionic diameter as Ca^2+^ (∼1.0 Å) and can readily permeate MCUcx when divalent ions are chelated from the pore entrance by EDTA (17, 18, 30). This uptake of Na^+^ into mitochondria can be quantified as a decrease in mitochondrial potential (ΔΨ) using a potentiometric dye such as TMRM. The assay is reproducible and robust, particularly when TMRM is used in ratiometric mode (17), enabling biophysical investigations of MCUcx regulation in intact mitochondrial preparations when ionic gradients across the IMM cannot be controlled (37, 46).

We developed an improved Na^+^ uptake assay by including the Ca^2+^-selective ionophore ionomycin (47, 48) as well as an excess of EGTA in the extramitochondrial buffer to chelate any residual Ca^2+^ as it is transported from the intermembrane space (IMS) and matrix across the mitochondrial membranes by the ionophore (in exchange for 2H^+^) (Fig. S1A) (48). We also excluded all other cations and permeant anions from the buffer, leaving only Tris, HEPES, EGTA, sucrose, and succinate to preserve mitochondrial respiration. Na^+^ and EDTA were added as bolus injections during experiments. Although this Na^+^-driven Δψ response (NDDR) assay does not directly measure Na^+^ uptake, instead using TMRM as a semi-quantitative indicator of Na^+^-driven changes in Δψ, it avoids the significant limitations of Na^+^-sensitive dyes: suboptimal *K*_d_ values (insensitivity to low Na^+^ concentrations); challenging UV excitation requirements; and complications with selectivity and calibration due to the presence of other ions in mitochondrial subcompartments. This comparative measure of mitochondrial depolarization therefore provides a robust means to assess regulation of MCUcx.

We applied NDDR to the relatively pure and abundant mitochondria that can be readily obtained from mouse liver by differential centrifugation. Interestingly, in the presence of ionomycin, both the initial rate and the degree of Na^+^-induced mitochondrial depolarization (measured ∼3 mins after bolus injections of Na^+^ and EDTA) were markedly increased compared to assays in the absence of ionomycin (Fig. 1A-C). Ionomycin produced an ∼13-fold reduction in the *K*_0.5_ for Na^+^-induced depolarization (365.6 ± 72.8 mM versus 27.2 ± 2.3 mM Na^+^) and an ∼12-fold increase in the initial depolarization rate at 150 mM Na^+^ (0.403 %depolarization/sec versus 4.71 %depolarization/sec) (Fig. 1B, C). We found similar effects of ionomycin in other model systems, including mitochondria isolated from mouse kidneys as well as permeabilized cells from several mouse and human cell lines (Fig. 1 and S1). Notably, mouse embryonic fibroblasts (MEFs) required ∼4 times less Na^+^ than human, HEK293, or HeLa cell lines or human induced-pluripotent stem cells (hiPSCs) to achieve a similar degree and rate of depolarization, with or without ionomycin (Fig. 1D-M, and S1D-M). This suggests there may be differences in the biophysical properties of MCUcx between immortalized cell lines. We also observed a mild hyperpolarization of mitochondria (∼4–5%) upon baseline application of ionomycin in both WT and MCUcx-deficient cells (Fig. 1D, F, I and K, and S1), as expected for exit of cations from the matrix. This agrees with a previous study, which found that the vast majority of ionomycin-induced transport was electroneutral (one Ca^2+^ exchanged for two H^+^) but that ∼10% of transport was electrogenic (49).

**Fig. 1.**
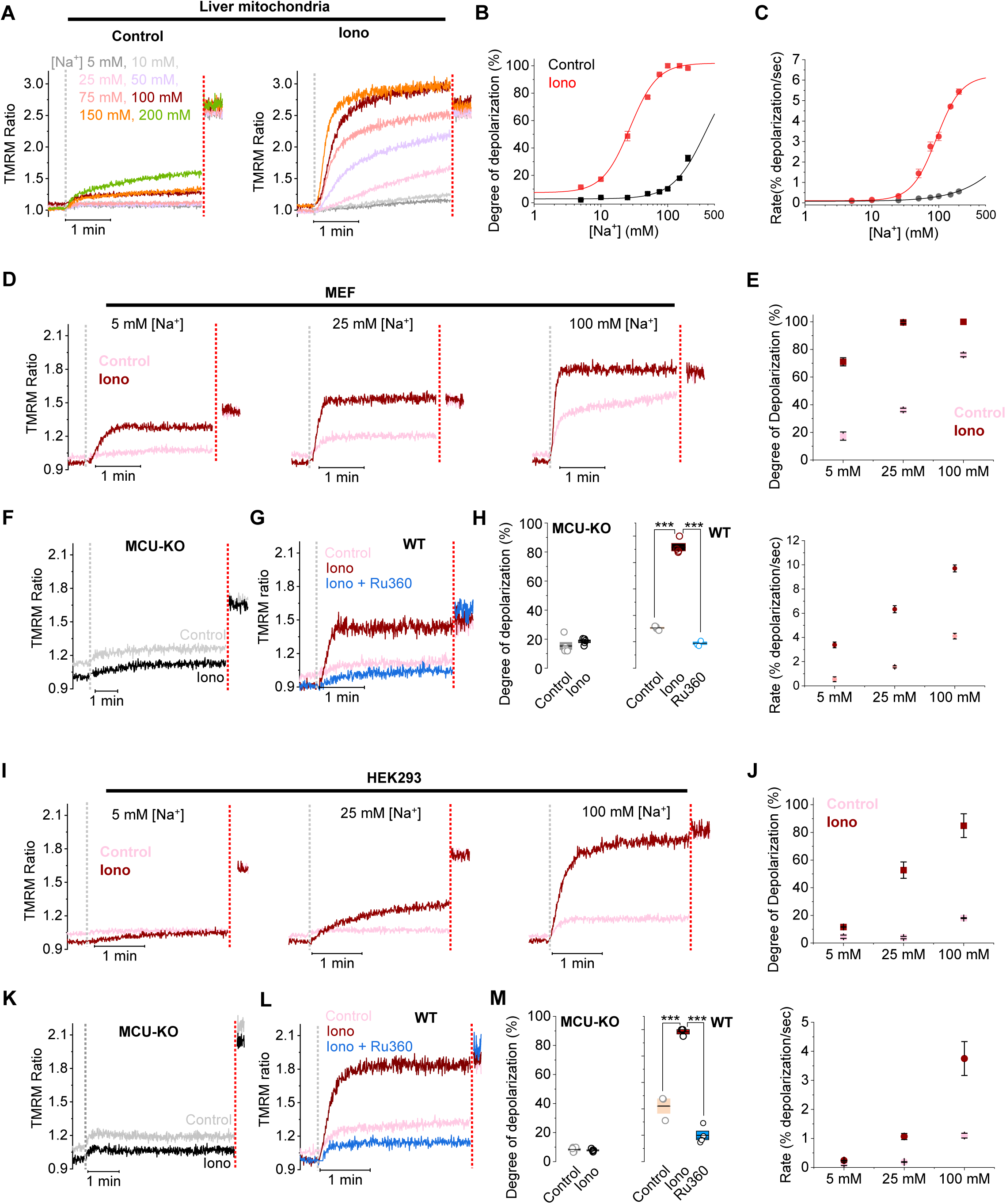
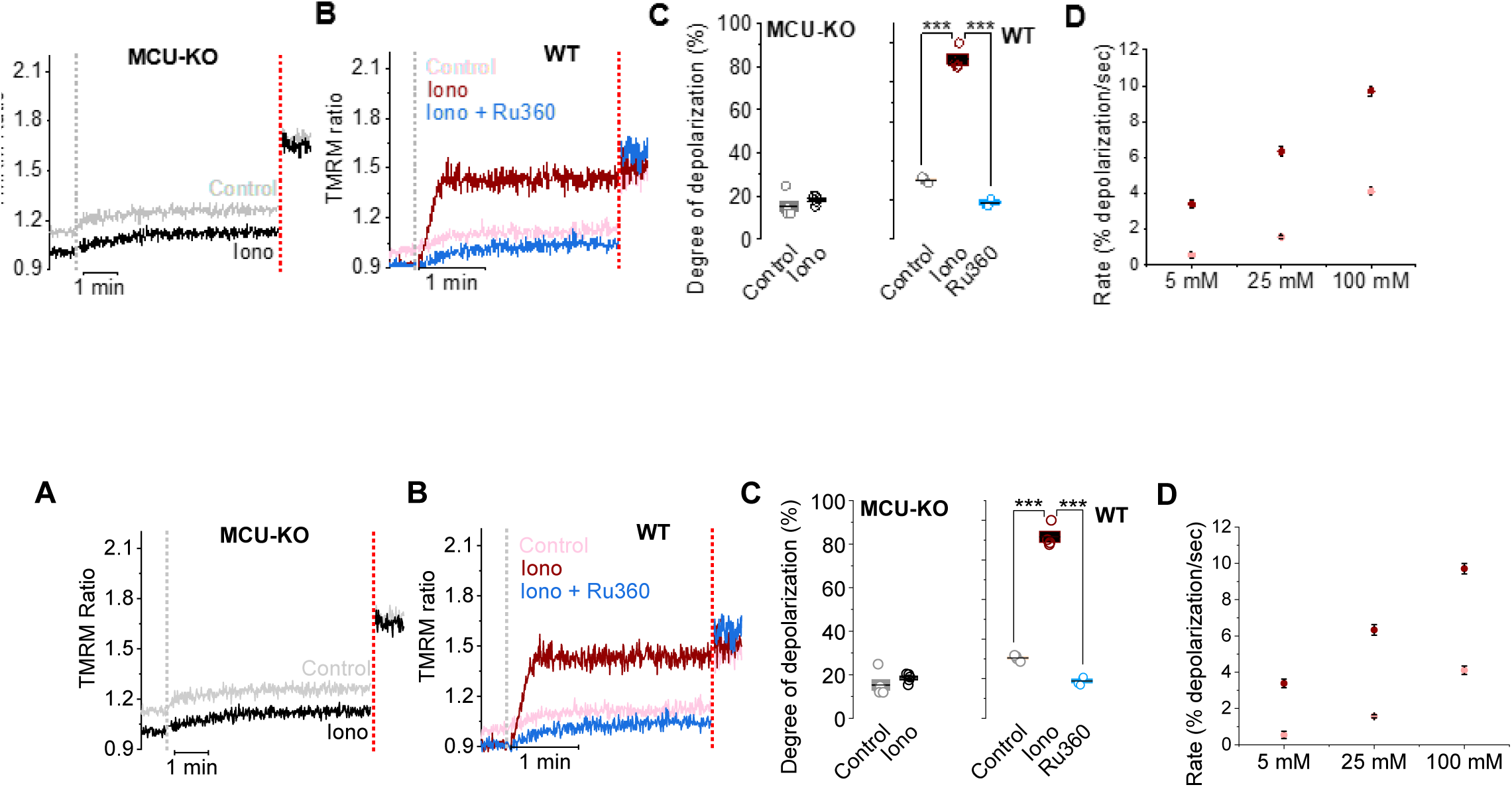
Ionomycin enables in situ measurements of MCUcx activity. (**A**) Representative time courses of Na^+^-uptake induced changes in Δψ (TMRM ratio) of isolated mouse liver mitochondria upon exposure to 10 mM EDTA and graded [Na^+^], in the absence (Control) or presence of 5 µM ionomycin (Iono). Grey dotted lines denote bolus injections of EDTA and Na⁺; red dotted lines mark FCCP addition. (**B and C**) Degree of depolarization (B), and initial depolarization rate (C) plotted as a function of [Na^+^]. Data is fitted to a Logistic equation. *K*_0.5_ for degree of depolarization is 27.2 ± 2.3 mM (*n* = 2.1 ± 0.2) for ionomycin and 365.6 ± 72.8 mM (*n* = 1.8 ± 0.3) for Control. (**D, F and G**) Time course of Na^+^-uptake induced Δψ changes in permeabilized Drp1-deficient mouse embryonic fibroblasts (MEF) for MCU KO (F), and cells with intact MCU (D and G) in the absence (Control) or presence of ionomycin (Iono). Acute inhibition of MCUcx with Ru360 abolished the ionomycin-enhanced response (G). (**E and H**) Degree of depolarization (E, upper panel and H) and the depolarization rate (E, lower panel) for MEF. (**I, K and L**) Time course of Na^+^-uptake induced Δψ changes in permeabilized HEK293 for MCU KO (K), and cells with intact MCU (I) in the absence (Control) or presence of ionomycin (Iono). Acute inhibition of MCUcx with Ru360 abolished the ionomycin-enhanced response (L). (**J and M**) Degree of depolarization (J, upper panel and M) and the depolarization rate (J, lower panel) for HEK293. Summary data presented as mean ± SEM; *\*\*\*P < 0.001*; two-sample t-test, two-tailed; n = 3-5.

To exclude the possibility that ionomycin has non-specific effects on mitochondrial potential, for example by acting on the OXPHOS complexes or allowing Na^+^ uptake via an MCU-independent route, we investigated Na^+^-induced mitochondrial depolarization in MCU and EMRE genetic knockouts (KOs). Na^+^ uptake was negligible and insensitive to ionomycin following MCU KO, EMRE KO, and MCU/EMRE double KO, in both MEFs (Fig. 1F, 1H, and S1H and I) and HEK293 cells (Fig. 1K, 1M and S1J and K), confirming that our assay measures Na^+^ uptake by MCUcx specifically and that ionomycin enhances the sensitivity of the assay by increasing the rate and extent of MCU-mediated Na^+^ uptake. We used 25 mM [Na^+^] for MEFs and 100 mM for HEK293 cells, as these concentrations maximized the ionomycin-induced response and fully utilized the dynamic range of the assay. Because the assay has high temporal resolution, we are able to observe dynamic modulation of TMRM ratio as Na^+^ influx proceeds (Fig. 1A) due to the effect of this cation flux on Δψ. We therefore used the initial rate of Δψ depolarization in our analyses, reflecting the early phase of Na^+^ entry when the driving force is relatively strong. Acute inhibition of MCUcx with Ru360 similarly suppressed Na^+^-induced depolarization in WT MEFs and HEK293 cells (Fig. 1G, H, L and M), further confirming that NDDR specifically reflects MCUcx-mediated ion flux. Together, these data show that NDDR can specifically quantify MCU function, with sensitivity and high temporal resolution, in a versatile range of systems with intact mitochondria.

### Matrix Ca^2+^ depletion enhances MCUcx-mediated ion flux

To determine whether and how matrix Ca^2+^ affects MCUcx function, we combined our NDDR assay with Ca^2+^-sensitive dyes and genetically-encoded indicators targeted to mitochondria (Fig. 2). Because isolated mitochondria and permeabilized cells were suspended in 2 mM EGTA, no extramitochondrial Ca^2+^ was detected upon application of ionomycin to mitochondria isolated from mouse liver (Fig. S2A). However, when EGTA was omitted from the suspension buffer, a clear Fura-8FF signal corresponding to ∼10 µM Ca^2+^ was detected, indicating that ionomycin caused significant amounts of Ca^2+^ to be released from these mitochondria (Fig. 2A). Regardless of EGTA presence at the beginning of the assay, mitochondria completely depolarized when a bolus of Na^+^ and EDTA was added to the suspension (Fig. 2A, lower panel, and S2A and B).

**Fig. 2.**
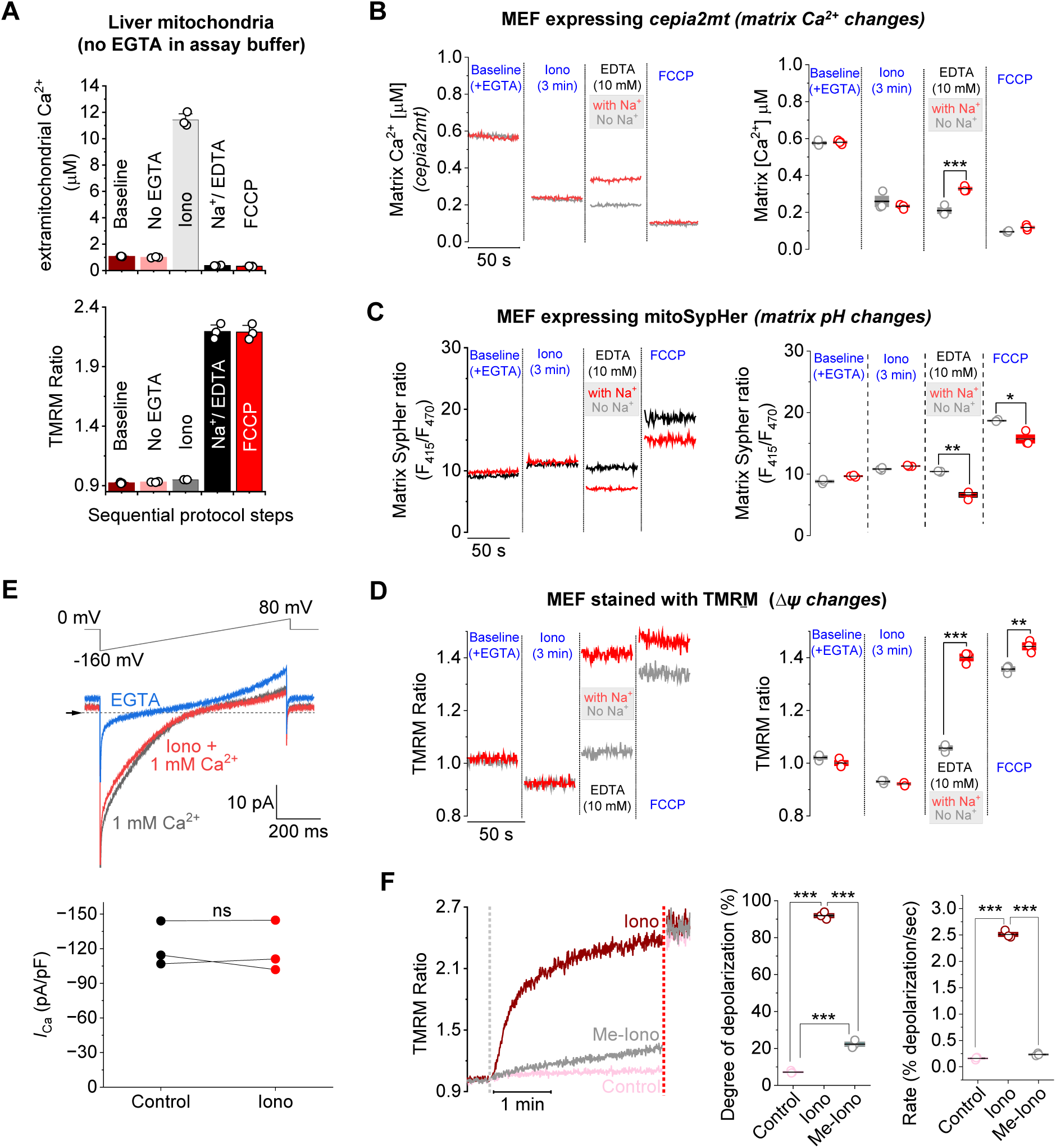
Matrix Ca^2+^ depletion enhances MCUcx-mediated ion flux. **A**) Extramitochondrial [Ca^2+^] measured at each step of the assay (upper panel) using isolated mouse liver mitochondria. EGTA is omitted from the assay buffer to permit detection of Ca^2+^ eleased from mitochondria following ionomycin treatment. Lower panel shows corresponding Δψ values from the same experiment. (**B**) Time course of matrix [Ca^2+^] in permeabilized MEFs tably expressing the mitochondria-targeted Ca^2+^ indicator, *cepia2mt*, at each step of the assay left panel). Experiments performed either in the absence (grey) or presence of Na^+^ (red). Right anel shows summary data for matrix [Ca^2+^]. (**C**) Time course of *mito-SypHer* signal, a matrix-argeted pH indicator, in permeabilized MEFs at each step of the assay (left panel). Experiments performed either in the absence (grey) or presence of Na^+^ (red). Right panel hows summary data of relative *mito-SypHer* signal changes (F415/F470). (**D**) Time course of Δψ in permeabilized MEFs at each step of the assay (left). Experiments performed either in the bsence (grey) or presence of Na^+^ (red). Right panel shows summary data of changes in Δψ. Summary data presented as mean ± SEM; \**P < 0.05, **P < 0.01, ***P < 0.001*; two-sample t-est, two-tailed; n = 3-5. (**E**) Representative MCUcx-mediated whole-IMM Ca^2+^ currents ecorded from isolated mouse liver mitoplasts in the presence of 1 mM Ca^2+^ before (black) and fter application of 5 µM ionomycin + 1 mM Ca^2+^ (red) (upper panel). Baseline currents (blue race) measured in HEPES/Tris solution. Voltage protocol shown above current traces. Lower anel shows Ca^2+^ current density measured at -156 mV. ns, not significant; Paired Sample t-est; n = 3. (**F**) Time course of Na^+^-uptake induced Δψ changes in MEFs in the absence Control) or presence of ionomycin (Iono), or in the presence of non-Ca^2+^-chelating methyl ster ionomycin (Me-Iono). Right panels show summary data for degree of depolarization and epolarization rate. Grey dotted line denotes bolus injection of EDTA and Na⁺; red dotted line marks FCCP addition. Summary data presented as mean ± SEM; \**P < 0.05, **P < 0.01, ***P < .001*; One-way ANOVA with Bonferroni post hoc test; n = 3.

We next examined changes in matrix Ca^2+^ levels upon ionomycin application using MEFs stably expressing the matrix-targeted genetically-encoded Ca^2+^ indicator *cepia2mt* (*K*_d_ ∼160 nM) (17, 50). Resting matrix Ca^2+^ in permeabilized cells suspended in assay buffer was ∼600 nM, which was reduced to ∼200 nM in the presence of ionomycin (Fig. 2B). Subsequent EDTA addition did not alter matrix [Ca^2+^] but addition of Na^+^ with EDTA (to trigger Na^+^ uptake) produced a modest transient increase in cepia2mt signal (∼50 nM equivalent change). Because this response was small and transient despite conditions designed to minimize Ca^2+^ availability, it is unlikely to reflect substantial or sustained matrix Ca^2+^ accumulation. Instead, the signal may arise from transient non-equilibrium changes associated with rapid Δψ changes (Fig. 2D), or other secondary effects on indicator behavior during Na⁺ influx. Thus, depletion of matrix [Ca^2+^] to ∼200 nM was sufficient to achieve ionomycin-mediated enhancement of Na^+^ influx. Addition of the protonophore FCCP at the end of the experiment further reduced matrix [Ca^2+^] due to complete depolarization of mitochondria (Fig. 2B and D). Moreover, an inhibitor of the mitochondrial Na^+^/Ca^2+^ exchanger, CGP37157, confirmed that both the initial rate and degree of depolarization in our NDDR assay are not influenced by this exchanger (Fig. S3). Together, these results demonstrate that the combination of EGTA, EDTA, and ionomycin promotes substantial net depletion of matrix Ca^2+^ under NDDR conditions.

To assess changes in matrix [H^+^] during the NDDR assay, we used MEFs stably expressing the mitochondria-targeted ratiometric pH indicator *mitoSypHer* (Fig. 2C). Application of ionomycin in the presence of EGTA modestly decreased the pH gradient across the IMM (acidifying the matrix) due to import of two H^+^ for every Ca^2+^ exported. Addition of EDTA restored matrix alkalinity back to baseline, likely by chelating divalent cations required to support continued ionophore activity. Addition of Na^+^ in the presence of EDTA produced a small decrease in the mito-SypHer ratio, indicating modest matrix alkalinization. This change coincided with marked Na^+^-induced mitochondrial depolarization measured by TMRM (Fig. 2D), suggesting that it reflects secondary adjustments in matrix proton balance during rapid mitochondrial depolarization.

To exclude the possibility that ionomycin is directly binding to MCUcx and acting as a ligand, we conducted mitoplast patch-clamp experiments, as Ca^2+^ uptake assays cannot be used to measure membrane protein function in the presence of a Ca^2+^ ionophore. We recorded MCU-mediated Ca^2+^ currents from individual mitoplasts isolated from mouse liver (17, 37) and confirmed that ionomycin has no effect on currents through MCUcx in the presence of 1 mM Ca^2+^ (Fig. 2E). We therefore hypothesized that removal of Ca^2+^ from the mitochondrial matrix by ionomycin has a direct effect on MCUcx function. We tested this by synthesizing a methyl ester derivative of ionomycin (Me-Iono) that can no longer bind Ca^2+^ with high affinity, allowing us to distinguish the Ca^2+^-dependent effects of ionomycin from any off-target or membrane-perturbing actions. Indeed, the methyl ester derivative elicited minimal changes in the initial rate or degree of mitochondrial depolarization in the NDDR assay, compared to unaltered ionomycin (Fig. 2F). Moreover, in contrast to ionomycin (Fig. 2A), the methyl ester form was unable to release Ca^2+^ from the mitochondrial subcompartments when applied under identical conditions (Fig. S4). These results strongly support the conclusion that Na^+^-driven Δψ responses in the presence of ionomycin are driven by depletion of matrix Ca^2+^ rather than nonspecific effects on the IMM. We thus conclude that matrix Ca^2+^ inhibits MCUcx-mediated ion influx via a novel mechanism, which we term “matrix Ca^2+^ inhibition” (MCI), that likely involves a Ca^2+^-binding or regulatory site in the matrix. The ease with which our NDDR assay can be combined with different tools also demonstrates the versatility of our approach.

### Electrophysiological analysis of MCUcx in mitoplasts and orthogonal controls for divalent ion-dependent regulation

To further evaluate the mechanistic basis of matrix Ca^2+^-dependent MCUcx regulation and address potential assay-specific confounders, we performed a series of complementary electrophysiological, pharmacological, and ionophore-based experiments (Fig. 3). Because previous electrophysiological studies (32, 33) proposed that matrix divalent ions directly suppress MCU activity, we first tested whether matrix-side Ca^2+^ alters MCU-mediated currents in whole-mitoplast recordings under conditions similar to those previously described. Addition of 400 nM Ca^2+^ to the matrix-facing pipette solution produced no inhibition of MCU current density in isolated mouse liver mitoplasts (Fig. 3A and B), despite robust matrix Ca^2+^-dependent inhibition observed in intact mitochondria using NDDR (Fig. 1A-C). These findings suggest that matrix Ca^2+^-dependent inhibition may depend on features of the intact mitochondrial environment that are not fully preserved in mitoplast preparations, including native matrix and intermembrane space milieu, membrane organization, and ion buffering.

**Fig. 3.**
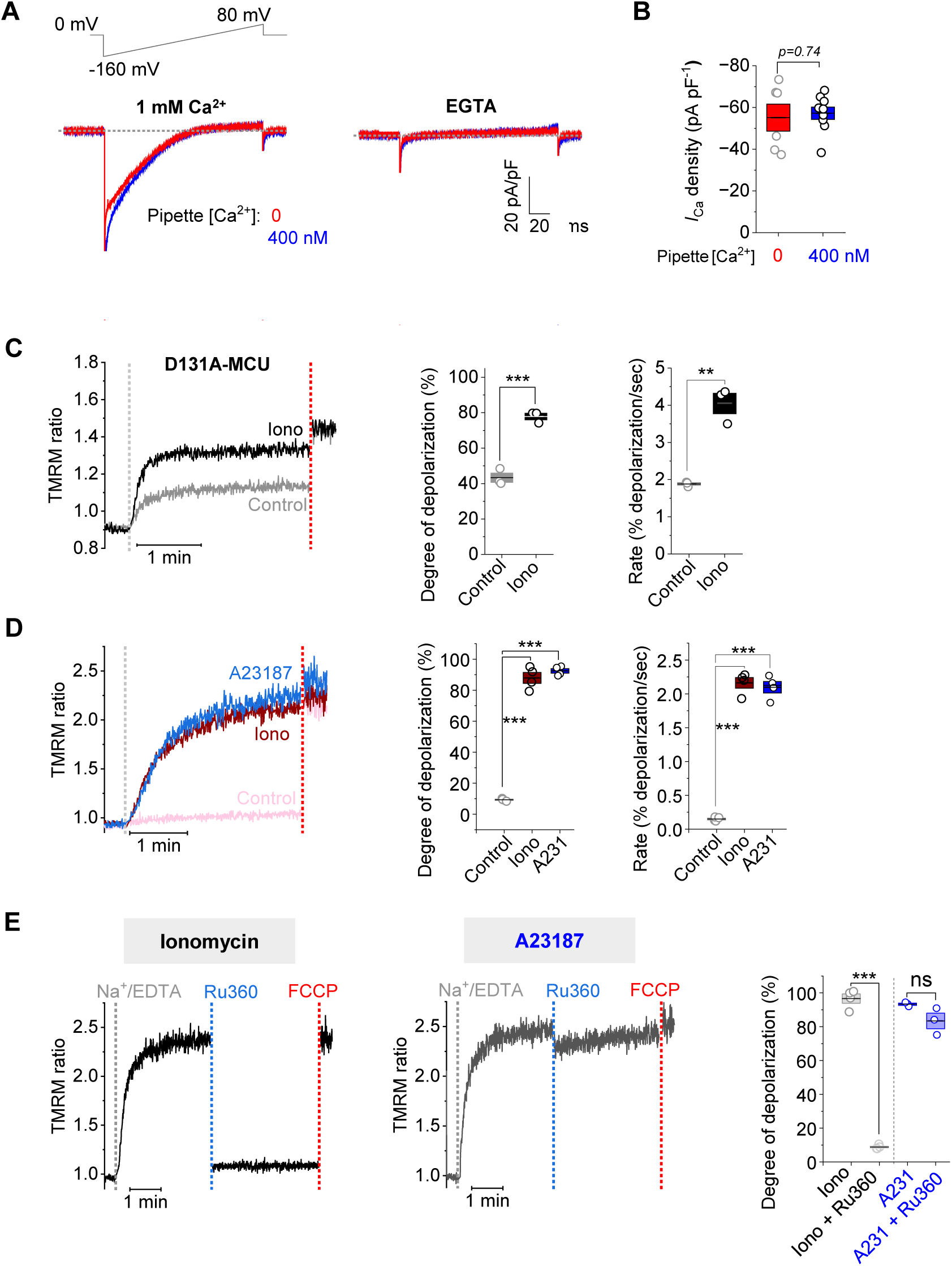
Electrophysiological analysis of MCUcx in mitoplasts and orthogonal controls for divalent ion-dependent regulation. **A**) Representative whole-mitoplast MCU currents (*I*_Ca_) from isolated mouse liver mitoplasts nduced by 1 mM bath Ca^2+^ (left) or baseline currents in the presence of bath EGTA (right), with ither nominally Ca^2+^-free (0) or 400 nM Ca^2+^ in the matrix-facing pipette solution. Voltage rotocol shown above traces. (**B**) Quantification of *I*_Ca_ density recorded at −156 mV under the ndicated pipette [Ca^2+^] conditions. Matrix-side 400 nM Ca^2+^ did not significantly alter MCU- mediated Ca^2+^ currents. n = 6-9 (**C**) Time course of Na^+^-uptake induced Δψ changes in MCU KO HEK293 cells expressing D131A MCU. Grey dotted lines denote bolus injections of EDTA nd Na⁺ [100 mM]; red dotted lines mark FCCP addition. Degree of depolarization (middle anel) and depolarization rate (right panel) of D131A MCU. D131A-MCU retained sensitivity to onomycin-mediated enhancement of Na^+-^induced depolarization. (**D**) Time course of Na^+^-ptake induced Δψ changes in isolated liver mitochondria comparing the effects of different onophores, relatively Ca^2+^-selective ionomycin, and Mg^2+^/Ca^2+^ non-selective A23187 (A231), ach 5 µM. Quantification of the degree of depolarization and initial rate of depolarization is hown on the right. (**E**) Time course of Na^+^-uptake induced Δψ changes in isolated liver mitochondria showing the effects of acute MCU inhibition by Ru360 in the presence of onomycin (left) or A23187 (right). FCCP was added at the end of each recording to fully issipate Δψ. Quantification of Ru360-mediated inhibition of Na^+^-induced depolarization is hown on the right. Acute Ru360 application restored Δψ in the presence of relatively Ca^2+^-elective ionomycin but not in the presence of divalent ion non-selective A23187. Grey dotted ine denotes bolus injection of EDTA and Na⁺; blue dotted line marks Ru360 addition; red otted line marks FCCP addition. Summary data presented as mean ± SEM; \**P < 0.05, **P < .01, ***P < 0.001*; two-sample t-test, two-tailed; n = 3-5.

We next examined an N-terminal acidic patch to which Mg^2+^ was bound in a structure of the isolated N-terminal fragment of MCU (residues 72-189) (34). This acidic domain bound divalent ions with millimolar affinity (*K*_d,Ca2+_ ∼2.3 mM, *K*_d,Mg2+_ ∼1.7 mM), causing monomerization of the N-terminal fragment. Intriguingly, neutralization or charge reversal of amino acids forming the divalent binding site (D131 and D147) decreased MCU activity in imaging assays (34) and rendered MCU insensitive to intra-matrix Ca^2+^-dependent inhibition in patch clamp recordings (33). We therefore utilized the same D131A MCU-expressing cell line as the above studies (33, 34) but observed intact sensitivity to ionomycin (Fig. 3C) arguing against a primary role for this site in the inhibitory mechanism observed in intact mitochondria and thus suggests that a different mechanism is responsible for MCI.

To determine whether the enhanced Na^+^ flux observed during NDDR was specific to Ca^2+^ or instead reflected a more general consequence of matrix divalent ion depletion including Mg2+, we compared ionomycin with the structurally distinct, divalent ion non-selective ionophore A23187 under identical experimental conditions. A23187 produced Δψ responses that initially resembled those observed with ionomycin (Fig. 3D), indicating that perturbation of matrix divalent ion homeostasis can enhance mitochondrial depolarization under NDDR conditions. However, when Na^+^/EDTA-induced depolarization was allowed to proceed to an apparent steady state before acute inhibition of MCUcx with Ru360, only ionomycin-treated mitochondria exhibited restoration of Δψ, whereas A23187-treated mitochondria remained largely insensitive to Ru360 (Fig. 3E). The ability of Ru360 to restore Δψ following ionomycin treatment demonstrates that mitochondria retained substantial respiratory capacity and that the observed depolarization primarily reflected ongoing MCUcx-mediated ion influx rather than irreversible dysfunction of the electron transport chain. In contrast, the lack of recovery following A23187 treatment is consistent with broader perturbation of divalent ion homeostasis by this non-selective ionophore. Together, these orthogonal experiments support the conclusion that ionomycin primarily reveals MCUcx-dependent regulation in intact mitochondria while highlighting the importance of Ca^2+^ specificity.

### Matrix Ca^2+^-dependent inhibition (MCI) is functionally distinct from MICU1-mediated gatekeeping

Because MICU1 gives rise to Ca^2+^-dependent regulation, specifically inhibition of MCUcx via a steric effect on the pore entrance during resting cytosolic [Ca^2+^] conditions (17, 25, 26, 51), we investigated whether MICU1 might be involved in MCI. To rule out the possibility that Ca^2+^ released by ionomycin binds to MICU1 and relieves channel inhibition, we utilized MICU1-KO cells. As expected, the absence of MICU1 significantly enhanced the rate and degree of depolarization before ionomycin application, regardless of cell type (MEFs, Fig. 4A-D; HEK293, S5A-D). Importantly, ionomycin further increased the rate and extent of depolarization in MICU1-KO cells, demonstrating that matrix Ca^2+^-dependent inhibition remains detectable in the absence of MICU1-mediated gatekeeping. The effect of ionomycin was particularly evident when [Na^+^] was lowered from 25 to 5 mM, as this limited mitochondrial depolarization and prevented saturation of the TMRM signal, thereby improving sensitivity by keeping the response within the assay’s dynamic range (Fig. 4A, B (lower panels) and D, and S5A, B (lower panels) and D).

**Fig. 4.**
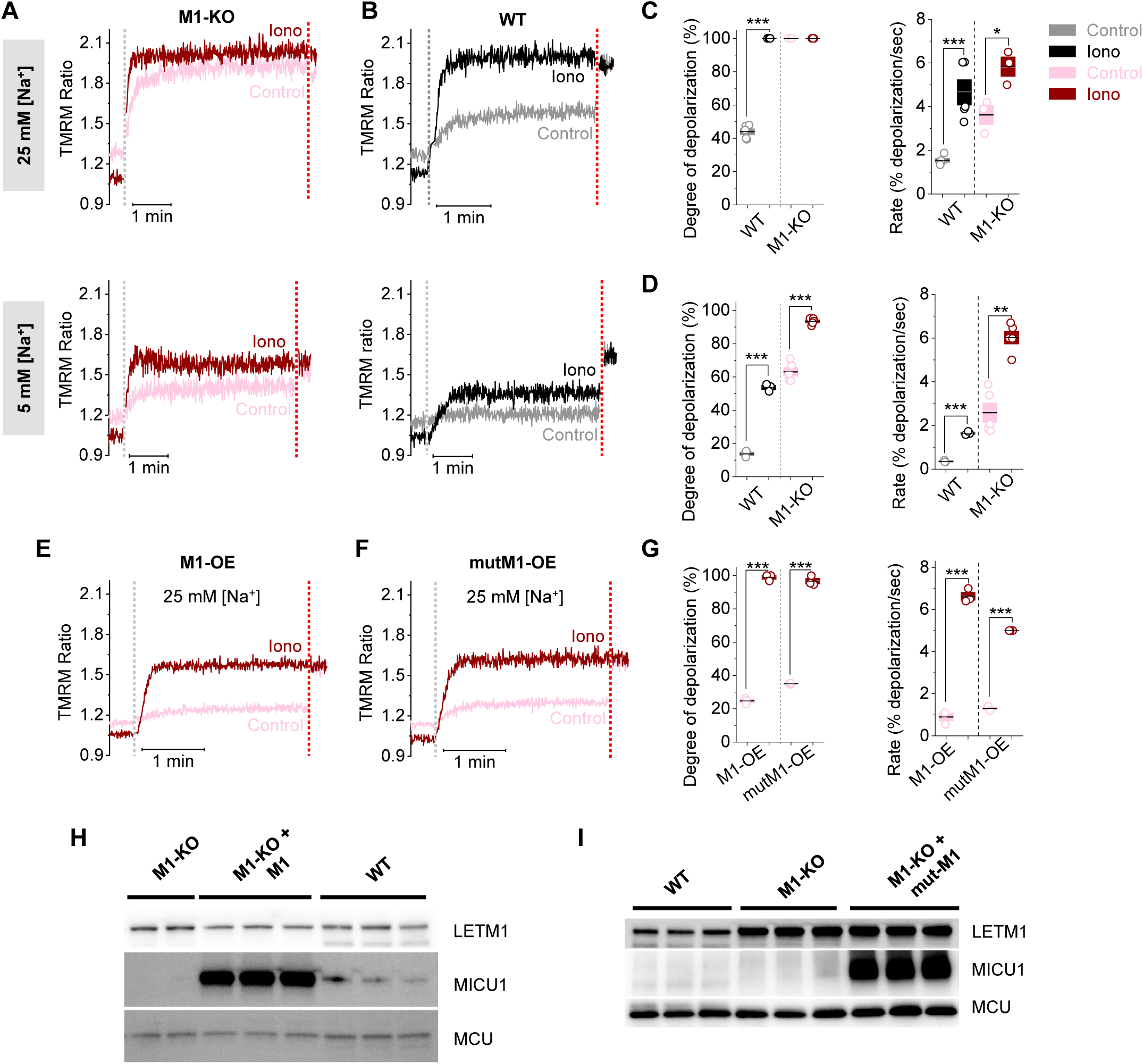
Matrix Ca^2+^-dependent inhibition is functionally distinct from MICU1-mediated gatekeeping. (**A**) Time course of Na^+^-uptake induced Δψ changes in MICU1-KO MEFs (M1-KO) following addition of 25 mM (upper) and 5 mM Na^+^ (lower) in the absence (Control) or presence of ionomycin (Iono). (**B**) For comparison, representative time courses for Na^+^-uptake induced Δψ changes in permeabilized WT cells shown under the respective conditions. (**C**) Degree of mitochondrial depolarization (left) and depolarization rate (right) of MICU1-KO and WT MEFs in 25 mM Na^+^. (**D**) Degree of mitochondrial depolarization (left) and depolarization rate (right) of MICU1-KO and WT MEFs in 5 mM Na^+^. (**E and F**) Time course of Na^+^-uptake induced Δψ changes in MICU1-KO MEFs stably expressing WT MICU1 (M1-OE, E) or EF hand mutated MICU1 (mutM1-OE, F) using 25 mM Na^+^ in the absence (Control) or presence of ionomycin (Iono). (**G**) Degree of depolarization and depolarization rate of cell lines in (E and F). (**H and I**) Western blot showing abundant WT MICU1 or EF hand mutated MICU1 protein in respective stable cell lines, MICU1-KO MEFs stably expressing WT MICU1 (M1-KO + M1, H) or EF hand mutated MICU1 (M1-KO + mut-M1, I). Grey dotted lines denote bolus injections of EDTA and Na⁺; red dotted lines mark FCCP addition. Summary data presented as mean ± SEM; *\*\*\*P < 0.001*; two-sample t-test, two-tailed; n = 3-5.

Interestingly, in MICU1-KO HEK293 cells, Na⁺-driven Δψ responses in the presence of ionomycin exhibited a distinct biphasic profile, characterized by an initial rapid depolarization peaking at ∼30% above control levels followed by a slower relaxation to the same steady state as controls (Fig. S5A (lower panel) and D). A qualitatively similar yet more subtle biphasic response was noticeable in MICU1-KO MEFs, where ionomycin predominantly enhanced the rate and extent of depolarization (Fig. 4A (lower panel) and D). These data suggest subtle cell type variations in MCUcx regulation or different ion homeostasis mechanisms. Regardless, our results show that, although MCI remains intact in MICU1-KO cells, there is a marked reduction in the [Na^+^] required for full mitochondrial depolarization. The increased sensitivity of MICU1-KO cells to Na^+^-induced depolarization indicates that MICU1 strongly influences the overall magnitude of matrix Ca^2+^-dependent regulation. However, because ionomycin-dependent enhancement of Na^+^ flux remains detectable, matrix-side inhibitory mechanisms are not completely abolished in the absence of MICU1.

The persistence of ionomycin-induced enhancement of Na^+^-driven depolarization in MICU1-KO cells suggested that MCI is not fully accounted for by MICU1-mediated gatekeeping alone. We therefore investigated whether increasing MICU1 expression could further suppress MCUcx-mediated Na+ flux and alter the ionomycin-dependent response by stably overexpressing WT MICU1 in MICU1-KO cells using lentiviral transduction, to avoid the mixed population of cells and therefore heterogenous measurements that can result from transient transfection (Fig. S5E and F). This stable cell line robustly expressed WT MICU1 protein, resulting in more than an order of magnitude greater expression than endogenous MICU1 in WT cells (Fig. 4H), (Fig. 4E and G) and almost completely suppressing Na^+^ influx in the absence of ionomycin. However, these cells still exhibited a pronounced acceleration in the rate and extent of depolarization upon ionomycin application (Fig. 4E and G). These findings indicate that even marked overexpression of MICU1 is insufficient to completely eliminate ionomycin-dependent enhancement of MCUcx-mediated ion flux. Thus, although MICU1 serves as a critical gatekeeper that sets the threshold for MCU activation, matrix Ca^2+^-dependent inhibitory mechanisms can still modulate channel behavior under these experimental conditions.

Although the buffer used contains both EGTA and EDTA, it is possible that Ca^2+^ released from the matrix by ionomycin may activate MICU1. We tested this in MICU1-KO cells stably overexpressing a mutant form of MICU1 with EF-hands that cannot bind Ca^2+^ (mutMICU1) and are therefore in a persistent apo state. Cells stably expressing mutMICU1 had more than an order of magnitude greater expression than endogenous MICU1 in WT cells (Fig. 4I). Because mutMICU1 is in a persistent apo-state, and results in marked inhibition of mitochondrial Ca^2+^ uptake (52–54), we expected minimal Na^+^ uptake in our NDDR assay. However, mutMICU1-overexpressing cells exhibited substantial Na^+^ uptake in the absence of ionomycin, with a kinetic profile similar to that in cells overexpressing WT MICU1 cells (Fig. 4F and G). We furthermore observed an increase in the initial rate and degree of mitochondrial depolarization in the presence of ionomycin, ruling out a requirement for Ca^2+^ binding to MICU1 in the ionomycin-induced enhancement of MCUcx-mediated ion flux, despite the ability of MICU1 to influence the magnitude of MCI. Together, these findings demonstrate that matrix Ca^2+^-dependent inhibition remains detectable in the absence of MICU1 and can be experimentally distinguished from MICU1-mediated gatekeeping using NDDR in intact mitochondria.

### EMRE and a juxtamembrane loop in the MCU subunit are essential for MCI

We next sought to identify the Ca^2+^ binding site responsible for MCI by focusing on the matrix domains of the MCU subunit. We first generated a large N-terminal truncation variant of MCU (Δ57-189 MCU) and additional point mutations of negatively charged residues (E229A and E287S). All these variants were functional and displayed intact MCI (Fig. 5A and B, and S6A-D), ruling out a major contribution of these sites to MCI.

**Fig. 5.**
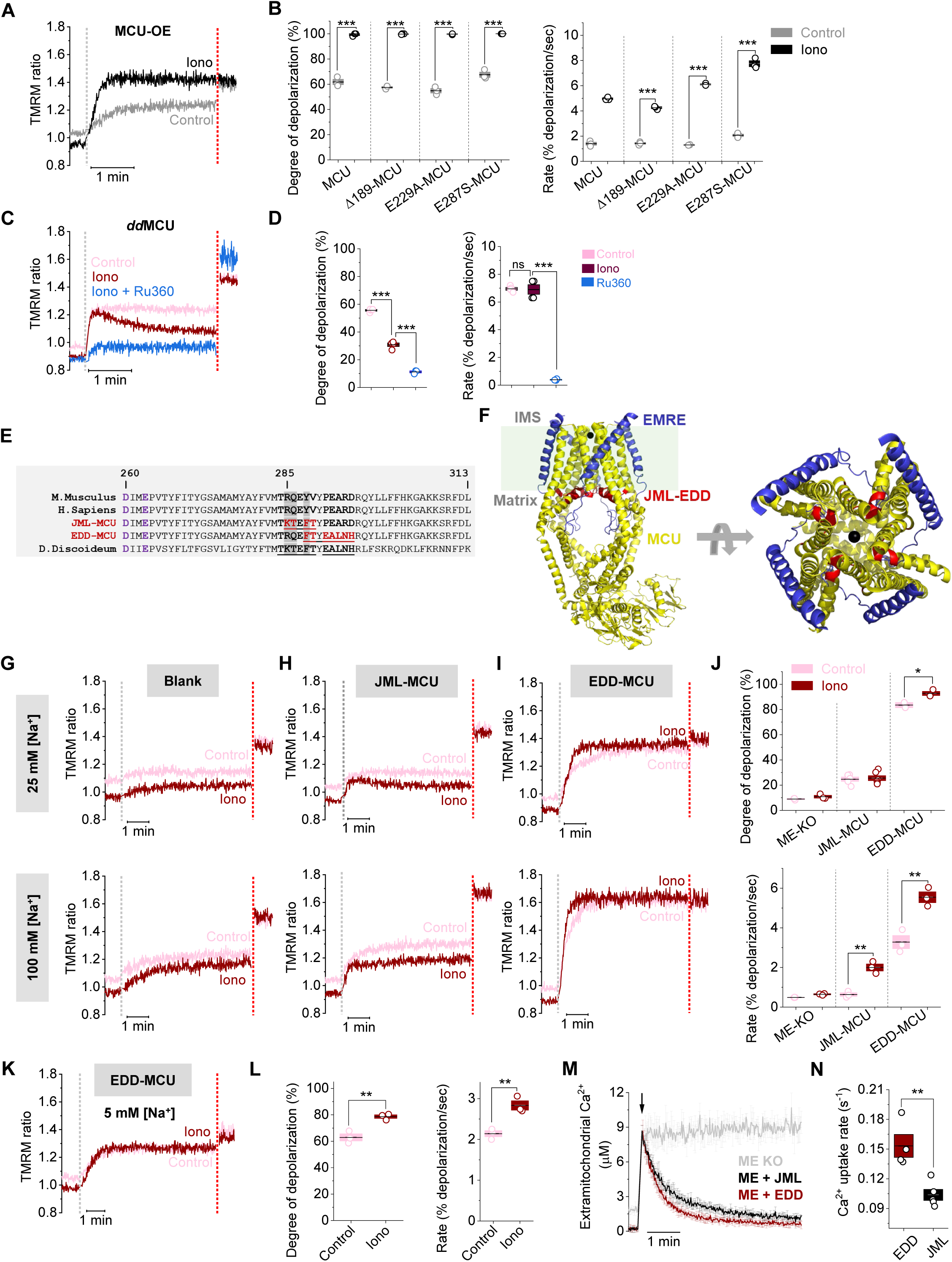
EMRE and a juxtamembrane loop in the MCU subunit are essential for MCI. (**A**) Time course of Na^+^-uptake induced Δψ changes in MCU-KO MEFs expressing WT MCU (MCU-OE). [Na^+^] is 25 mM. (**B**) Degree of mitochondrial depolarization (left) and depolarization rate (right) for respective MCU variants expressed in MCU-KO MEFs. (**C**) Time course of Na^+^-uptake induced Δψ changes in MCU-KO HEK293 expressing *dd*MCU. [Na^+^] is 100 mM. (**D**) Degree of mitochondrial depolarization (left) and depolarization rate (right) of *dd*MCU. (**E**) Sequence alignment of MCU homologues. Numbers indicate position of mMCU residues. (**F**) Structural model of the MCUcx (PDB 6O58, left) showing the JML-EDD region (red) within the tetrameric MCU subunit (yellow). Right panel shows a view from the top (cytosolic-side). EMRE is shown in blue and Ca^2+^ in black. (**G**-**I**) Time course of Na^+^-uptake induced Δψ changes in MCU/EMRE double KO MEFs (Blank, G) or stably expressing JML-MCU (H) or EDD-MCU (I). [Na^+^] is 25 mM (upper panels) or 100 mM (lower panels). (**J**) Degree of mitochondrial depolarization (upper) and depolarization rate (lower) in 25 mM Na^+^ for respective EMRE-independent MCU variants. (**K**) Time course of Na^+^-uptake induced Δψ changes in MCU/EMRE double KO MEFs expressing EDD-MCU. [Na^+^] is 5 mM. (**L**) Degree of mitochondrial depolarization (left) and depolarization rate (right) in 5 mM Na^+^ for EDD-MCU. (**M**) Averaged traces show mitochondrial Ca^2+^-uptake kinetics in permeabilized MCU/EMRE double KO MEFs (ME-KO) or cells expressing JML-MCU (ME + JML) or EDD-MCU (ME + EDD). Arrow marks the time of bolus Ca^2+^ injection. (**N**) Quantification of mitochondrial Ca^2+^ uptake rates. Data presented as mean ± SEM; \**P < 0.05, **P < 0.01, ***P < 0.001*; two-sample t-test, two-tailed; n = 3-5.

*Dictyostelium discoideum*, a member of the phylum Amoebozoa, expresses a MCU homolog (*dd*MCU) that functions independently of EMRE (36). A juxtamembrane loop (JML) region on the matrix-facing side of ddMCU is known to facilitate this independence (19). To investigate whether MCI is conserved in EMRE-independent MCU homologs, we used MCU/EMRE double-KO cells stably expressing ddMCU. Interestingly, the initial rate and magnitude of Na^+^ uptake were largely similar in the absence and presence of ionomycin, in contrast to the robust ionomycin-dependent enhancement observed with mammalian MCU (Fig. 5C and D). In addition, in the presence of ionomycin, Na^+^ uptake peaked at the same value as the control, but subsequently relaxed to a steady state of approximately half the peak magnitude. Because Ru360 inhibited the ddMCU-mediated response (Fig. 5C and D), these results demonstrate that ddMCU remains capable of conducting Na^+^ under NDDR conditions and that the altered phenotype does not simply reflect loss of channel activity. Instead, the biphasic response indicates that matrix Ca^2+^ regulation is fundamentally altered in ddMCU relative to mammalian MCU, despite conservation of the Ca^2+^-binding residues that form the selectivity filter (Fig. 5E, purple color residues). These findings suggest that the mammalian MCI phenotype cannot be explained by changes in Ca^2+^ occupancy of the conserved selectivity filter residues and instead involves additional structural determinants outside the pore. Together, these findings suggest that EMRE-dependent regulation and/or structural elements within the JML region are important for coupling matrix Ca^2+^ depletion to MCUcx activity.

To distinguish between these two hypotheses, we employed two chimeric constructs incorporating short overlapping regions from *dd*MCU into mammalian MCU, each of which should confer independence from EMRE (Fig. 5E and F). One construct contained a short substitution (residues 285–289) to replace the native JML with that of *dd*MCU (JML-MCU) (19). The other included substitution of the native EMRE dependence domain (EDD; residues 288–95 that putatively interact with EMRE) (31) with the equivalent region in *dd*MCU (EDD-MCU). These variants were stably expressed in MCU/EMRE double KO cells (Fig. 5G-N). Although Na⁺-induced depolarization was observed in cells expressing EDD-MCU (Fig. 5I and J, and S6E), the enhancement in initial rate and magnitude with ionomycin was markedly less than WT MCU, suggesting that MCI is present but highly diminished. This effect was also observed when the experiment was repeated using 5 mM Na^+^ to maintain the Δψ response within the dynamic range of TMRM (Fig. 5K and L). In contrast, ionomycin had almost no effect on the degree of mitochondrial depolarization in JML-MCU-expressing cells, accompanied by only an acceleration of the depolarization rate. Overall, JML-MCU responses closely resembled those observed in MCU/EMRE double-knockout cells (Fig. 5G, H, J and S6E).

Together, these findings indicate that EMRE is a key determinant of MCI, with the JML region serving as a critical interface through which this regulatory signal is conveyed to the MCU pore. While disruption of the JML region strongly impairs MCI, the marked attenuation observed upon EMRE perturbation demonstrates that EMRE actively contributes to the efficiency and dynamic range of this inhibitory mechanism. Furthermore, our data raises the possibility that EMRE (perhaps via its interaction with EDD) and the JML region act together, either as part of a Ca^2+^-responsive module or through cooperative structural interactions, to integrate Ca^2+^-dependent signals to control MCUcx gating. Previous work (19) showing that substitution of the *dd*MCU JML is sufficient to confer EMRE-independent activity supports the idea that this region serves as a key regulatory interface linking EMRE to channel function. Notably, conventional mitochondrial Ca^2+^ uptake assays revealed robust Ca^2+^ uptake by both JML-MCU and EDD-MCU variants (Fig. 5M and N), indicating good expression and full functionality. However, these assays detected only subtle differences between the two variants, highlighting the importance of the superior sensitivity and temporal resolution of our NDDR assay (Fig. 5H-J) for resolving MCUcx functional differences.

### EMRE enhances MCUcx ion permeation and channel throughput

The findings above highlight that EMRE plays an important functional role beyond its necessity for MCU channel formation and MICU1 tethering in higher eukaryotes(19–21). However, its exact contribution is unclear, particularly given the EMRE-independent MCU channels in some species (35, 36). To investigate the physiological role of EMRE, we sought to engineer a minimally-altered, EMRE-independent MCU variant with intact EMRE interaction regions. By comparing the protein sequences of WT MCU, EDD-MCU, and JML-MCU (Fig. 5E), we identified R285K and Q286T as the only substitutions distinguishing JML-MCU from both WT and EDD-MCU, and subsequently constructed an R285K/Q286T (RQ-MCU) mutant. This variant preserved crucial interactions between EMRE and MCU, including those with transmembrane domains 1 and 2 as well as the matrix-facing EDD region (19, 21, 31), and exhibited robust expression and functionality in both MCU-KO and MCU/EMRE double KO cell lines (Fig. 6A-F).

**Fig. 6.**
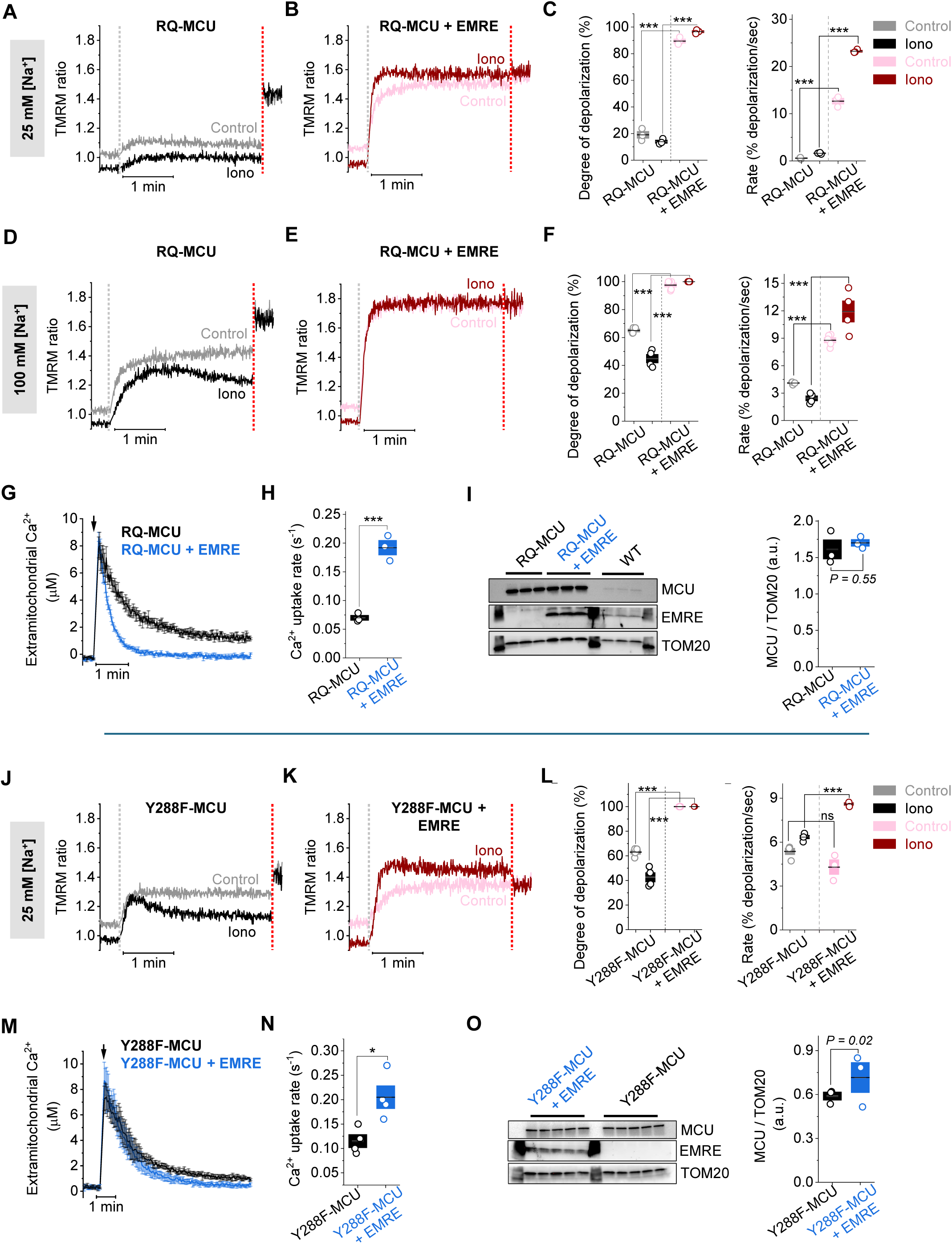
EMRE enhances MCU ion permeation and channel throughput. (**A and B**) Time course of Na^+^-uptake induced Δψ changes in MEFs stably expressing R285K/Q286T MCU in either MCU/EMRE double KO background (RQ-MCU, A) or MCU-KO background (RQ-MCU + EMRE, B). Grey dotted line denotes bolus injection of EDTA and Na⁺ [25 mM]; red dotted line marks FCCP addition. (**C**) Degree of mitochondrial depolarization (left) and depolarization rate (right) for RQ-MCU expressing lines. (**D**) Traces showing mitochondrial Ca^2+^-uptake kinetics in MEFs stably expressing RQ-MCU. Arrow marks time of bolus Ca^2+^ injection. (**E**) Quantification of mitochondrial Ca^2+^ uptake rate in respective cell lines. (**F**) Western blot showing RQ-MCU protein expression in MEFs stably expressing R285K/Q286T MCU in either MCU/EMRE double KO background (RQ-MCU) or MCU-KO background (RQ-MCU + EMRE). Right panel shows quantification of RQ-MCU band intensity. (**G, H and I**) Representative time courses of Na^+^-uptake induced Δψ changes in MEFs stably expressing Y288F-MCU in either MCU/EMRE double KO background (Y288F-MCU, G) or MCU-KO background (Y288F-MCU + EMRE, H), and quantification of kinetic parameters (I). (**J and K**) Representative traces showing mitochondrial Ca^2+^-uptake kinetics in MEFs stably expressing Y288F-MCU (J), and quantification of mitochondrial Ca^2+^ uptake kinetics in the respective cell lines (K). (**L**) Western blot showing Y288F-MCU protein expression and its quantification in the respective MEFs. Data is presented as mean ± SEM; *\*\*\*P < 0.001*; One-way ANOVA with Bonferroni post hoc test or two-sample t-test (two-tailed); n = 3-5.

In the presence of EMRE, RQ-MCU-expressing cells exhibited near-complete depolarization in 25 mM Na^+^ regardless of ionomycin presence (Fig. 6B and C). In contrast, in the absence of EMRE, RQ-MCU-expressing cells exhibited NDDR responses in 25 mM Na^+^ that resembled those of MCU-KO cells (Fig. 6A and C, and Fig. 1F) suggesting markedly reduced ion permeation or an intrinsically unstable conductive state under these conditions. We confirmed robust RQ-MCU functionality in the absence of EMRE using higher [Na^+^] (100 mM, Fig. 6D-F), which led to ∼50% mitochondrial depolarization in the absence of ionomycin. Interestingly, ionomycin elicited an initial depolarization peak that was comparable to control, followed by ∼40% repolarization over ∼3 mins (Fig. 6D), resembling the response of *dd*MCU (Fig. 5C). This was in contrast to the response in the presence of EMRE, when RQ-MCU exhibited complete depolarization at both 25 mM (Fig. 6B and C) and 100 mM Na^+^ (Fig. 6E and F), regardless of ionomycin, and did not display a biphasic response. Time course analysis revealed that EMRE increased the initial depolarization rate as well as the degree of depolarization (Fig. 6C and 6F), suggesting that it facilitates more efficient permeation through the pore. This marked facilitation was observed despite Western blot analysis revealing similar RQ-MCU expression in EMRE-expressing cells compared to EMRE-deficient cells (Fig. 6I), ruling out a contribution of protein expression and underscoring the direct functional impact of EMRE.

To further corroborate the functional role of EMRE revealed by the RQ-MCU variant, we tested another MCU variant, Y288F, where a relatively conserved mutation was shown to enable MCU opening without EMRE (55). Interestingly, Y288 residue is also a part of the JML region (Fig. 5E). In EMRE-deficient (MCU/EMRE double KO) cells, Y288F-MCU elicited partial (∼60%) depolarization in response to 25 mM Na^+^ (Fig. 6J and L). Addition of ionomycin produced a biphasic response (Fig. 6J), closely resembling the behavior observed for *dd*MCU, and RQ-MCU in the absence of EMRE. In contrast, expression of Y288F-MCU in the presence of EMRE (MCU-KO cells) (Fig. 6K) resulted in nearly complete mitochondrial depolarization at the same Na^+^ concentration, regardless of ionomycin. Ionomycin addition further accelerated the rate of depolarization, consistent with EMRE enhancing both channel throughput and matrix Ca^2+^-dependent regulation (Fig. 6K and L). Interestingly, EMRE-expressing Y288F-MCU cells exhibit only marginally faster Ca^2+^ uptake than EMRE-deficient cells (Fig. 6M and N) compared to the obvious functional differences in the NDDR assay. While Western blot analysis shows slightly higher Y288F-MCU levels in EMRE-expressing cells, (Fig. 6O), this minor difference in protein abundance complicates the interpretation of Ca^2+^ uptake kinetics, whereas EMRE’s functional contribution remains distinct and obvious in the NDDR assay. Importantly, EMRE-independent variants remained functional in both NDDR and conventional Ca^2+^ uptake assays, indicating that EMRE is not absolutely required for ion permeation per se, but instead promotes a robustly conductive state capable of supporting high levels of ion flux.

Taken together, these results indicate that EMRE promotes a highly conductive state of MCUcx, thereby enhancing ion permeation and overall channel throughput. EMRE-independent MCU variants reveal that, in the absence of EMRE, MCU occupies a low-throughput or weakly conductive state that nevertheless retains measurable ion permeation, explaining why EMRE is required for robust activity. The high temporal resolution and sensitivity of our NDDR assay provide clear and direct evidence to support this role, in contrast to Ca^2+^-uptake experiments, which confirmed that EMRE-expressing RQ-MCU or Y288F-MCU cells exhibit faster and greater Ca^2+^ uptake than EMRE-deficient cells but with more equivocal evidence.

## DISCUSSION

The ability to dynamically regulate mitochondrial Ca^2+^ uptake via MCUcx is crucial to balance the metabolic benefits of mitochondrial Ca^2+^ signaling with protection against Ca^2+^ overload and cell injury. Here, we define the mechanistic basis of EMRE’s essential role in MCUcx regulation by identifying two fundamental and mechanistically distinct roles of EMRE. Using MCU variants that are EMRE-independent yet retain intact EMRE-interaction regions, we show that EMRE is required for robust channel activity and additionally fine-tunes Ca^2+^ permeation under physiological conditions.

These advances have been made possible by the development and rigorous validation of a versatile assay, NDDR, that can specifically measure MCUcx-mediated ion uptake with high sensitivity and temporal resolution in intact mitochondrial preparations. NDDR decouples MCUcx-mediated ion permeation from the confounding influences of Ca^2+^ ions, buffering systems, and organellar cross-talk, while preserving native protein interactions, regulatory mechanisms, and the milieu of mitochondrial subcompartments. The advantages of NDDR over conventional Ca^2+^ uptake measurements are due to its quantitative rigor, specifically, temporal resolution and sensitivity. This allowed us to distinguish subtle biophysical differences between MCU variants, which are often obscured in cumulative Ca^2+^ uptake assays, and identify partial loss-of-function, mosaic expression, and regulatory tuning of channel activity. Given the crucial role of MCUcx in neuromuscular, cardiac, immune, and metabolic disorders, we believe the advantages of NDDR will catalyze elucidation of novel regulatory mechanisms, annotation of the MCUcx interactome (22), and screening of both disease-linked variants and pharmacological modulators of MCUcx.

An important consideration is that NDDR probes MCUcx function under non-equilibrium conditions that differ fundamentally from both conventional Ca^2+^ uptake assays and electrophysiological recordings. By combining ionophore-mediated matrix divalent depletion with Na^+^-driven flux measurements in intact mitochondria, NDDR preserves the native architecture of the organelle, including matrix and IMS milieu, membrane organization, and protein interactions. Consistent with this distinction, the matrix Ca^2+^-dependent inhibitory phenotype described here was readily detected in intact mitochondria but not in mitoplast recordings performed under comparable ionic conditions, as also reported previously (17). Although the basis for this discrepancy remains unresolved, our findings suggest that matrix-side regulation of MCUcx may depend on physiological features of the intact organelle that are not fully preserved following mitoplast preparation. Importantly, NDDR should not be viewed as a direct surrogate for physiological mitochondrial Ca^2+^ uptake. Rather, it is a mechanistic tool that experimentally separates ion permeation from the confounding effects of Ca^2+^- dependent regulation, buffering or transport itself. The ability to independently manipulate matrix divalent ions while monitoring MCUcx-mediated flux enables regulatory mechanisms that would otherwise be obscured during conventional Ca^2+^ uptake measurements. Future studies will be required to determine the exact molecular basis of MCI, define its relationship to previously proposed matrix regulatory mechanisms, and establish the extent to which it contributes to mitochondrial Ca^2+^ homeostasis in vivo.

Using this approach, we discovered MCI – a negative feedback mechanism distinct from inhibition by MICU1 in low cytosolic Ca^2+^ whereby elevations in matrix [Ca^2+^] suppress MCUcx-mediated ion flux – extending the foundational work by Heaton and Nicholls on the use of Ca^2+^ ionophores for studying mitochondrial Ca^2+^ transport (1976) (48). Our current work incorporates genetic and structural analyses of MCUcx and reveals that matrix Ca^2+^ can directly inhibit MCUcx activity, which remains intact even in the absence of MICU1. Previous electrophysiological studies proposed that MICU1 contributes to matrix Ca^2+^-dependent inhibition of MCU activity (32, 33), and our findings are broadly consistent with a role for MICU1 in this process, although the magnitude and manifestation of the inhibitory phenotype differ across genotypes and experimental systems. We furthermore show that MCI is dependent on the JML motif in the MCU subunit and that interaction with EMRE is required for optimal Ca^2+^-dependent gating. Importantly, MCI remained intact in cells expressing the D131A MCU mutant previously implicated in matrix divalent ion sensing. Together with the lack of inhibition observed in mitoplast recordings under comparable matrix Ca^2+^ conditions, these findings suggest that the inhibitory mechanism revealed by NDDR is not fully explained by previously proposed models and may involve additional structural elements of the intact uniporter complex. While Ca^2+^ occupancy of the selectivity filter could contribute to reduced Na^+^ permeation under some conditions, this passive occupancy model does not readily explain the coordinated dependence of the response on EMRE, the JML region, and MICU1. Thus, our data suggests that matrix Ca^2+^ modulates the conductive state of the MCU complex through regulatory interactions, rather than acting solely as a pore blocker. Physiologically, MCI may couple MCUcx activity to the metabolic state of the cell, permitting dynamic tuning based on local Ca^2+^ levels, redox signals, and Ca^2+^ buffering capacity (34, 41, 56). It may also serve to desensitize the uniporter during pathological Ca^2+^ elevations, preventing mitochondrial overload and preserving Δψ (57). Intriguingly, local nanodomains of Ca^2+^, generated by an open MCU pore, could themselves trigger MCI, establishing a self-limiting feedback loop that is tightly integrated with mitochondrial signaling. These feedback properties may also intersect with the rapid mode of uptake (RaM), a transient high-conductance state of MCUcx that mediates fast Ca^2+^ influx during cytosolic Ca^2+^ spikes (41, 58). When pulses occur too frequently to allow a full reset of RaM, matrix Ca^2+^ may accumulate and engage MCI, terminating RaM-like activity and limiting flux duration to preserve mitochondrial homeostasis. The molecular identity of the matrix divalent sensing mechanism remains unresolved. Our mutational analysis excludes several candidate sites within MCU and instead points toward cooperative contributions from the JML region and EMRE, providing a framework for future structural and mechanistic studies.

Our work also extends the functional repertoire of EMRE beyond its established role as a MICU1 tether and enabler of a functional MCU channel. By engineering MCU variants that bind to EMRE but remain functional in its absence, we were able to uncouple functional pore formation from regulatory modulation. Targeted mutations confined to the JML of MCU (at R285, Q286 and Y288), while preserving known EMRE interaction interfaces, including the transmembrane domains and the EDD (19, 31, 55), demonstrate that uniporter opening involves substantial conformational rearrangements within the JML-EDD region (Fig. 5F). This strategy further revealed that EMRE enhances MCUcx throughput, likely by favoring a highly conductive pore conformation and increasing the efficiency of ion permeation. Although the presence of EMRE increased the rate of Ca^2+^ uptake, the functional effect was more clearly resolved using the NDDR assay, due to its superior sensitivity to kinetic differences. The larger EMRE-dependent effects observed under Na^+^ compared with Ca^2+^ conditions are consistent with differences in basal *P*_o_ between these ions (17). Because *P*_o_ is intrinsically higher in Ca^2+^ than in Na^+^, EMRE-mediated increases in *P*_o_ produce a larger relative effect under Na^+^conditions, where baseline channel opening is more limited. The biophysical advantages of EMRE were evident in both steady-state and kinetic measurements, reinforcing a dual role for this key subunit in pore assembly and functional optimization. Although some species of fungi and Amoebozoa can produce functional MCU channels in the absence of EMRE (31, 35, 36), its acquisition may reflect an evolutionary adaptation toward more sophisticated regulatory control and enhanced ion flux efficiency, particularly during brief cytosolic Ca^2+^ signaling events.

## ACKNOWLEDGMENTS

We would like to thank Dr. David Nicholls (Buck Institute for Research on Aging, Novato, CA) and Dr. W. Jonathan Lederer (Center for Biomedical Engineering and Technology, University of Maryland School of Medicine, Baltimore) for insightful comments on the manuscript, and all members of the Garg laboratory for helpful discussions. We would like to thank Anu Sunkara for help with sample preparation for a few experiments. We would like to thank the University of Maryland School of Medicine Flow Cytometry Core-Baltimore, Maryland for the FACS services. We thank Drs. J. Kevin Foskett and Horia Vais (University of Pennsylvania, PA) for sharing the D131A-MCU cell line.

We kindly acknowledge the support from the National Institutes of Health (R01GM145806 [V.G.]), Maryland Stem Cell Research Fund (MSCRF- 96651 [V.G.]), University of Maryland Baltimore (Startup funds [V.G.]), American Heart Association (Postdoctoral award 24POST1241582 [A.K.]), and National Institute of Arthritis and Musculoskeletal and Skin Diseases (Training Program in Muscle Biology Grant T32 AR007592-27 [D.M.N.]). The content is solely the responsibility of the authors and does not necessarily represent the official views of the National Institutes of Health or any funding agency.

## AUTHOR CONTRIBUTIONS

A.K. and V.G. conceptualized the project, designed the experimental plan, analyzed the results, and co-wrote the manuscript. A.K. performed all the experiments and made the figures. V.G. supervised the project. A.K. and D.M.N. together designed and refined the assay. N.D. and J.P.Y.K. synthesized the methyl ester derivative of ionomycin. D.D. and S.H.K. assisted with molecular biology work. All authors contributed to the discussion of the results, editing of the manuscript and approve the final manuscript.

## DATA AND MATERIALS AVAILABILITY

All data needed to evaluate the conclusions in the paper are present in the paper and/or the Supplementary Materials. All the plasmids used in the paper will be deposited to Addgene and/or available from the corresponding author.

## MATERIALS AND METHODS

### Animals

All experiments involving wild-type mice (C57BL/6N) were obtained from the University of Maryland Baltimore (UMB) animal husbandry. These mice were maintained on a standard rodent chow diet and water under 12-hour light and dark cycles. Animal experiments were performed with male mice (2–5 months old) according to procedures approved by the University of Maryland Institutional Animal Care and Use Committee (IACUC) and in full compliance with the National Institutes of Health (NIH) guidelines for the care and use of laboratory animals.

### Cell culture

All cell lines (HEK293, MEFs, HeLa) were cultured in high-glucose Dulbecco’s modified Eagle’s medium (DMEM) supplemented with 10% fetal bovine serum (FBS), 100 U/mL penicillin, 100 U/mL streptomycin, and 0.1 mg/mL normocin at 37°C in a humidified incubator with 5% CO_2_. All cell lines were free of mycoplasma as determined by a PCR-based detection method. D131A-MCU expressing HEK293 cells (33, 34) were obtained from Drs. J. Kevin Foskett and Horia Vais (University of Pennsylvania, PA).

### Lentiviral packaging and stable cell line generation

HEK293T cells were cultured in 10 cm dishes at 70% confluency. Third-generation lentiviral plasmids mix encoding pMDL (Gag/Pol), pVSVG (vesicular stomatitis virus glycoprotein), pREV, and gene of interest with a selection marker (EGFP or mCherry) were used to transfect HEK293T cells with 25 kDa PEI. Plasmid vectors were either obtained from Vector Builder, Inc. (Chicago, IL, USA) or constructed in-house, with sequences verified by Plasmidsaurus whole-vector sequencing. Forty-eight and 72 hours post-transfection, lentivirus-containing supernatant was harvested, aliquoted, and stored at −80°C for further use. These lentiviral particles were used to make the stable cell line. Recombinant cDNA-expressing stable cells were generated by lentiviral transduction followed by selection using multiple rounds of fluorescence-activated cell sorting (FACS).

### Single-point or multiple-site mutagenesis

Site-directed mutagenesis was performed using either the Quikchange site-directed mutagenesis kit (ThermoFisher Sci. cat# 200518) or Q5 site-directed mutagenesis kit (New England Biolabs, cat# E0554S). In all site-directed construct preparations, the PCR and subsequent steps were performed as recommended by the kit manufacturer. Following transformation and plating, DNA from single colonies was screened using Sanger sequencing to confirm the presence of the desired mutation. Final validation of all constructs was performed using Plasmidsaurus whole-vector sequencing.

### Isolation of mouse liver and kidney mitochondria

Liver or kidney mitochondria were isolated from adult (8-10 weeks) mice using a protocol previously described (46). Briefly, mice were euthanized with CO_2_ asphyxiation using a CO_2_ gas cylinder, followed by cervical dislocation. The liver or kidney were excised and chopped into small pieces using scissors and rinsed to remove blood. Mitochondria were extracted by homogenizing the tissue in ice-cold initial media (IM) containing 260 mM sucrose, 20 mM HEPES, 2 mM EGTA, and 0.1% bovine serum albumin (BSA) (pH adjusted to 7.2 with Trizma base) using an ice-chilled Wheaton glass grinder with eight slow strokes of a Teflon pestle rotating at 600 rpm. The homogenate was centrifuged in a 15 mL Falcon tube at 700 × g for 6 minutes at 4°C to remove unlysed cells, followed by centrifugation at 3,200 × g for 10 minutes to collect the crude mitochondrial pellet. The pellet was washed once in ice-cold IM and centrifuged again at 3,200 × g for 10 minutes to obtain purified liver mitochondria.

### Na^+^-driven Δψ response (NDDR) assay

#### Permeabilized cells

All NDDR assays were performed using a BMG LABTECH CLARIOstar plate reader. Tetramethylrhodamine, Methyl Ester, Perchlorate (TMRM, ThermoFisher Scientific cat# T668) was used to monitor the mitochondrial potential (Δψ) response. Cells from three dishes were trypsinized, washed with PBS, and resuspended in a Ca^2+^-free wash buffer (260 mM sucrose, 20 mM HEPES, 2 mM EGTA), then stored at 4°C. Just before the assay, 1.5×10^6^ cells were permeabilized and mitochondria were energized by preincubating in Ca^2+^-free recording buffer (260 mM sucrose, 20 mM HEPES, 2 mM EGTA, 500 nM TMRM, 5 mM succinate, and 0.005% digitonin) with or without ionomycin (5 µM) for 3 minutes. The cell suspension was then loaded in a 96-well black plate (GreinerBio-One, cat# 650209).

TMRM fluorescence was recorded in the ratiometric mode (F_540_/F_575_), with 540-8 nm and 575-8 nm excitation and collection of emission at 619-30 nm. Following baseline acquisition, a bolus of NaCl with 10 mM EDTA was injected to trigger Na^+^ influx and a change in ΔΨ. At the end of each assay, 10 µM FCCP was added to completely dissipate ΔΨ. Summary data for the degree of depolarization is presented as a percentage of total dynamic range (baseline to FCCP or max depolarization), using the following equation:

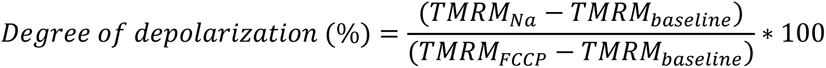

The initial rate of depolarization (calculated as % depolarization/sec, responses normalized to the full dynamic range of the TMRM signal) was calculated using Origin 2022b as a linear fit of the time-dependent change in Δψ response between the first 5-10 seconds following Na^+^/EDTA addition.

#### Isolated mouse tissue mitochondria

A similar protocol as above was used for mitochondria isolated from mouse tissues. Briefly, mouse liver mitochondria were isolated and resuspended in Ca^2+^-free wash buffer (260 mM sucrose, 20 mM HEPES, and 2 mM EGTA). Protein concentration was measured using BCA kit (ThermoFisher Scientific cat# 23227). Mitochondria (500 µg/mL) were energized in the recording buffer (260 mM sucrose, 20 mM HEPES, 1 mM EGTA, 500 nM TMRM, and 5 mM succinate) with or without ionomycin (5 µM) for 3 minutes. Ru360 (1 µM) was used to inhibit MCUcx-mediated ion flux. The mitochondrial suspension was then transferred to a well of a 96-well plate, and recordings were performed as described above.

### Mitochondrial Ca^2+^ uptake assay

Mitochondrial Ca^2+^ uptake experiments were carried out using a BMG LABTECH CLARIOstar plate reader. Extramitochondrial Ca^2+^ levels were monitored using a ratiometric, Ca^2+^-sensitive dye, Fura-8FF pentapotassium salt (AAT bioquest, cat# 20621, K_d_ ≈ 10 µM) or Fura-8 pentapotassium salt (AAT bioquest, cat# 21057, K_d_ ≈ 260 nM). Briefly, 1.5×10^6^ cells were suspended in Ca^2+^-free recording buffer (260 mM sucrose, 20 mM HEPES, 500 nm TMRM, 5 mM succinate, and 1 µM Fura-8FF) and incubated for 3 minutes at 4°C. Cells were then loaded in a 96 well black plate and a bolus of Ca^2+^ stock solution was injected. Fluorescence was recorded in the ratiometric mode (F_354_/F_415_) using excitation wavelengths of 354-8 nm and 415-8 nm, with emission at 524-8 nm. Calibration of Fura-8FF for [Ca^2+^] quantification was performed using this equation:

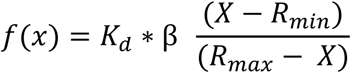

Where *X* represents the fluorescence intensity ratio, F_354_/F_415_ (F_354_=354-8 and F_415_=415-8 nm), which are the fluorescence detection wavelengths for the Ca^2+^-bound and Ca^2+^-free indicator, respectively. β is the ratio of F_min_ (at 0 Ca^2+^) to F_max_ (dye saturated with Ca^2+^) at 415-8 nm. *K*_d_ is the indicator’s Ca^2+^ dissociation constant. R_min_ is the fluorescence intensity ratio, F_354_/F_415_, at 0 Ca^2+^ and R_max_ is the fluorescence intensity ratio, F_354_/F_415_, at saturating Ca^2+^ concentration. Similarly the Fura8 or Fura8FF was used for the calcium release experiment from matrix.

The rate of Ca^2+^ uptake was calculated using Origin 2022b as a linear fit of the time-dependent change in extramitochondrial [Ca^2+^] during the first 5-10 seconds after bolus injection of Ca^2+^.

### Measurement of matrix [Ca^2+^]

Mitochondrial matrix Ca^2+^ levels were measured using MEFs stably expressing a genetically-encoded Ca^2+^ indicator, *cepia2mt* (*K*_d_ ∼160 nM), targeted to the mitochondrial matrix (17, 50). On the day of the experiment, cells were trypsinized and suspended in a Ca^2+^-free wash buffer (260 mM sucrose, 20 mM HEPES, 2 mM EGTA). Experiments with permeabilized cells were performed using 1.5 x 10^6^ cells preincubated for 3 minutes in a recording buffer (260 mM sucrose, 20 mM HEPES, 1 mM EGTA, 5 mM succinate, and 0.005% digitonin). Baseline fluorescence was recorded in a 96-well black plate by excitation at 470-15 nm and collection of emission at 515-20 nm. To assess ionomycin’s effect, cells were incubated with or without (control) 5 µM ionomycin for 3 minutes, and the fluorescence signal was recorded. Subsequently. EDTA was injected with or without Na^+^ and the resulting signal was measured. At the end of the experiment, FCCP was added, and *cepia2mt* fluorescence was recorded.

### Measurement of matrix pH

Mitochondrial matrix pH was measured using MEFs stably expressing a mitochondria-targeted ratiometric pH indicator, *mitoSypHer* (*59*). For the experiments, cells were trypsinized, and 1.5 x 10^6^ cells were permeabilized in recording buffer (260 mM sucrose, 20 mM HEPES, 1 mM EGTA, 5 mM succinate, and 0.005% digitonin) for 3 minutes. With excitation of *mitoSypHer* at 415-20 nm and 470-8 nm, fluorescence emission was acquired at 525/50 nm. After recording baseline, cells were incubated with or without 5 µM ionomycin for 3 minutes, and *mitoSypHer* fluorescence was measured. Fluorescence signal was recorded after injection of EDTA with or without Na^+^. At the end of the experiment, FCCP was added, and fluorescence was recorded. Data was plotted as the ratio of fluorescence intensities at F_415_/F_470_.

### Patch clamp recording of MCUcx currents

For the patch-clamp experiments, mitoplasts were prepared from isolated liver mitochondria, and whole-mitoplast currents were measured as described (46). Mitoplasts (mitochondria without outer membrane) were prepared using the Thermo Electron Corporation French press. Briefly, mitochondria were suspended for 5 minutes in hypertonic mannitol solution containing 140 mM sucrose, 440 mM D-mannitol, 10 mM HEPES, and 1 mM EGTA, and pH was adjusted to 7.2 with Tris base. After incubation, mitochondria were subjected to French press at ∼1100 psi. Mitoplasts were collected via centrifugation at 3800x g for 15 minutes and resuspended in hypertonic KCl solution (750 mM KCl, 100 mM HEPES, 1 mM EGTA, and pH adjusted to pH 7.2 with KOH) and stored at 4°C for further use for ∼4 hours. For the patch-clamp experiment, 20 µL of liver mitoplast suspension was added to 450 µL of KCl storing solution (150 mM KCl,10 mM HEPES, 1 mM EGTA, and pH adjusted to 7.0 with Trizma base) and plated on 5 mm coverslips pretreated with gelatin.

Giga-ohm seal of mitoplasts was formed in the bath solution containing 150 mM KCl, 10 mM HEPES, and 1 mM EGTA, pH 7.2 (adjusted with KOH). Voltage steps of 300–600 mV for 2–8 ms were applied to rupture the IMM and obtain the whole-mitoplast configuration. Typically, pipettes had a resistance of 25-35 MΩ, and the access resistance was 35–65 MΩ. The membrane capacitances of mitoplasts range from 0.2 to 0.8 pF.

For figure 2E, Na based pipette solution (PS) (110 mM Na-gluconate, 40 mM HEPES, 10 mM EGTA, and 2 mM MgCl_2_, pH 7.0 with NaOH, tonicity ∼350 mmol/kg) was used. For figure 3A, the TMA-based PS (130 mM TMA, 120 mM HEPES, 10 mM glutathione, and 2 mM MgCl_2_, pH adjusted to 7.0 with *D*-gluconic acid, tonicity adjusted to ∼346 mmol/kg with sucrose) was used with either 0 or 400 nM Ca^2+^.

### Preparation of ionomycin methyl ester

For the preparation of the methyl ester (Me-Iono), the procedure of Suzuki et al. 1987 (60) was used with minor modifications. Ionomycin (25 mg, 35 μmol, in 2.5 mL of ethanolic solution, Cayman Chemical, Ann Arbor, MI) was dried under vacuum to yield a yellow oil in a 4-mL reaction vial with a Teflon-silicone lined screw cap. To the vial, dry benzene (2 mL) and a small Teflon-coated stir bar were added; the mixture was stirred to give a solution. Two stock solutions were made: 1) iodomethane (22 μL, 350 μmol) dissolved in 500 μL dry benzene, and 2) 1,8-diazabicyclo(5.4.0)undec-7-ene (DBU, 53 μL, 350 μmol) dissolved in 500 μL dry benzene. 50 μL of each stock solution was added to the stirred ionomycin solution. The vial was purged with dry argon, sealed with Parafilm, and the solution was stirred for 4 days at room temperature in the dark. Thereafter, the solvent was removed by rotary evaporation. The resulting yellow oil was dissolved in CHCl3 and purified by flash chromatography on a prepacked silica gel column (elution with an ethyl acetate: CHCl3 gradient, 40% – 70% over 10 minutes; absorbance monitoring at 270 nm and 300 nm). The main peak was collected, and solvents were removed under vacuum to yield ionomycin methyl ester (19.3 mg, 26.7 μmol, 76% yield).

### Western blotting

Protein lysates were prepared as described in the previous manuscript (17). Total protein was extracted from the cells using ice-cold lysis media, radioimmunoprecipitation assay (RIPA) buffer (1% IGEPAL, 0.1% sodium dodecyl sulfate, 0.5% sodium deoxycholate, 150 mM NaCl, 1 mM EDTA, 50 mM Tris-HCl (pH 7.4), and a cocktail of protease inhibitors (Millipore Sigma, cat# 11836170001). Lysates were centrifuged at 14,000 × g for 10 minutes at 4°C, and protein concentrations were determined using the BCA assay.

Equal amounts of protein (15–20 μg) were resolved on 4–20% SDS-PAGE gels (Bio-Rad, cat# 4561095) and transferred onto PVDF membrane (Merck Millipore, cat# IPVH00010). Membranes were blocked and immunoblotted using the following primary antibodies: anti-MCU (Sigma, HPA016480, 1:2000), anti-MICU1 (Cell Signaling Technology, 12524S, 1:2000), anti-EMRE (Santa Cruz, sc-86337), anti-TOM20 (Santa Cruz, sc11415), and anti-LETM1 (ProteinTech, 16024-1-AP).

Anti-MICU1 antibody produced a non-specific band near its monomeric molecular weight (∼50 kDa), so samples were prepared in Laemmli buffer without β-mercaptoethanol to detect MICU1 homo- or heterodimers (∼100 kDa). Following primary antibody incubation, the membranes were washed and then incubated with appropriate HRP-conjugated secondary antibodies. The protein bands were visualized and imaged using an enhanced chemiluminescence detection system (ThermoFisher Sci. cat# 34580). ImageJ 1.54p was used for the quantitative analysis of protein expression levels.

### Materials

The following material were purchased: Ionomycin, 96% (ThermoFisher Sci., cat# j62448.MCR), NaCl (5M NaCl, Sigma cat# S5150-1L), EDTA (500 mM, Millipore, cat# 324504-500 mL), CaCl_2_ (Honeywell, cat# 21114-1L), Carbonylcyanide-p-trifluoromethoxyphenylhydrazone (FCCP, Sigma cat# C2920), Fura8FF^TM^ pentapotassium salt (AAT bioquest, cat# 20621), Fura8^TM^ pentapotassium salt (AAT bioquest, cat# 21057), Tetramethylrhodamine Methyl Ester, Perchlorate (TMRM, ThermoFisher Sci., cat# T668), Digitonin (5% Digitonin, ThermoFisher Sci. cat# BN20061), A23187 (Sigma, cat# C7522).

The following cell culture media supplies were purchased: Dulbecco’s Modified Eagle’s Medium (DMEM, high glucose: 4 mM L-glutamine, 4500 mg/L glucose, sodium pyruvate Cytiva Hyclone, cat# Sh30243.01), Fetal bovine serum (FBS, R&D systems cat# S12450), Phosphate buffered saline (PBS pH 7.4, Quality Biological, cat# 114-058-101), Trypsin-EDTA (0.25%, Gibco, cat# 25200072), Penicillin and streptomycin(100x, Corning, cat# 45000-653).

### Statistical analysis

Each experiment was performed using samples from 3-5 different cell culture dishes. Summary data are presented as mean ± standard error of the mean (SEM). The specific statistical tests used are indicated in the corresponding figure legends. Statistical analysis was completed in R (version 4.4.1) or Origin 2022b. *P* < 0.05 was considered statistically significant. Significance levels are indicated as follows: \**P < 0.05, **P < 0.01, ***P < 0.001*.

## SUPPLEMENTAL INFORMATION

Figures S1–S6.

