## Supplementary Information for "Mechanistic basis of EMRE’s essential role in the regulation of mitochondrial calcium uniporter complex"

**Figure S1.**

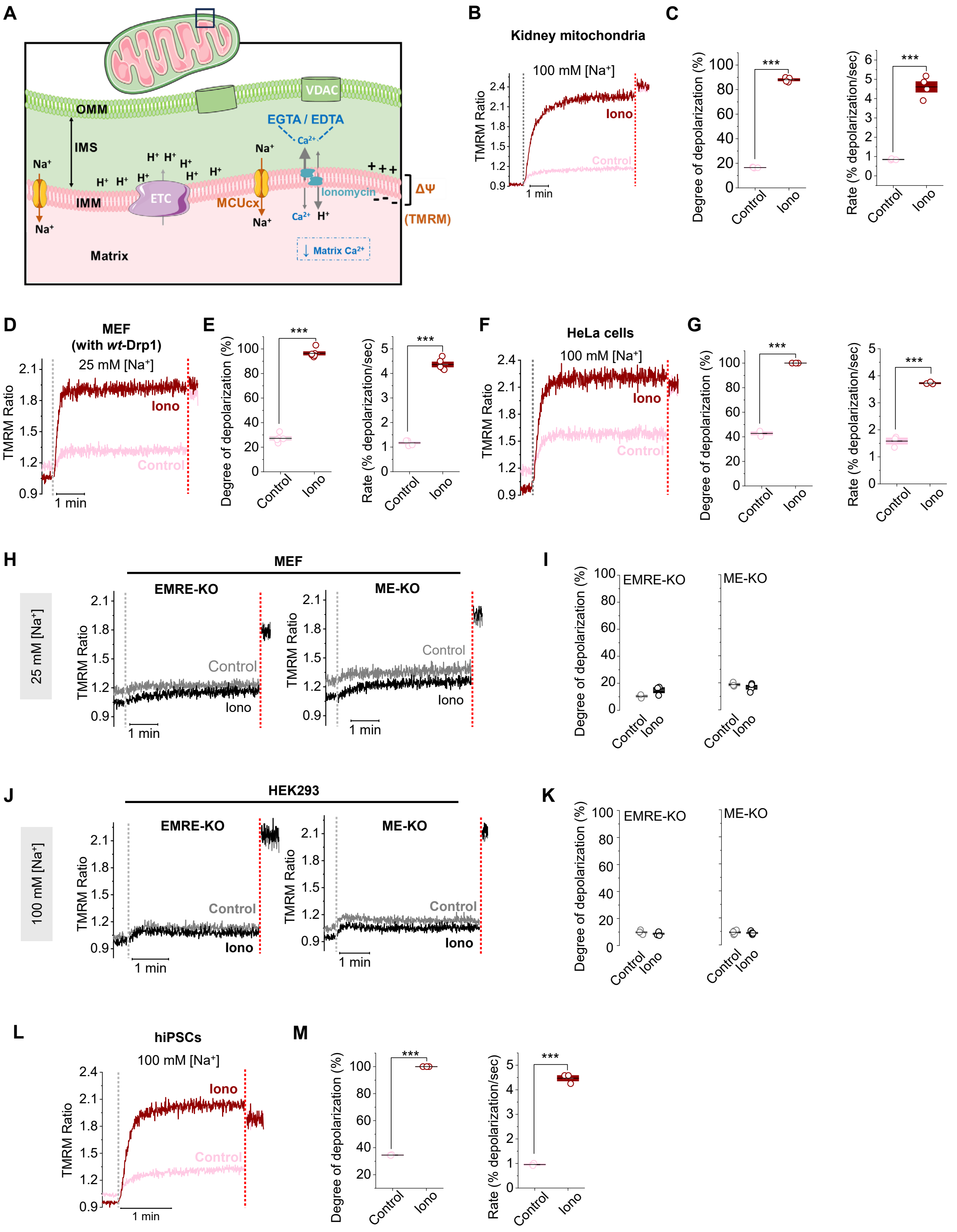
A
OMM
IMS
IMM
Matrix
VDAC
EGTA / EDTA
Ionomycin
ETC
MCUcx
ΔΨ
(TMRM)
↓ Matrix Ca2+
B
Kidney mitochondria
100 mM [Na+]
TMRM Ratio
Iono
Control
1 min
C
Degree of depolarization (%)
Rate (% depolarization/sec)
Control
Iono
***
D
MEF
(with wt-Drp1)
25 mM [Na+]
E
F
HeLa cells
100 mM [Na+]
G
H
MEF
EMRE-KO
ME-KO
25 mM [Na+]
I
J
HEK293
100 mM [Na+]
K
L
hiPSCs
100 mM [Na+]
M

**Fig. S1. Ionomycin enables in situ measurements of MCUcx activity.**

(**A**) Schematic illustrating the rationale for using ionomycin (**I**ono) in the NDDR assay. **I**onomycin in combination with EGTA/EDTA removes $Ca^{2+}$ from mitochondrial subcompartments, particularly the matrix, which is otherwise inaccessible to chelators. (**B**) Time-course of $Na^+$-uptake induced $\Delta\psi$ changes of isolated mouse kidney mitochondria using 100 mM [$Na^+$] in the absence (Control) or presence of ionomycin (**I**ono). Grey dotted lines denote bolus injection of EDTA and $Na^+$; red dotted lines mark FCCP addition. (**C**) Degree of depolarization (left) and the depolarization rate (right) for kidney mitochondria. (**D**) Time-course of $Na^+$-uptake induced $\Delta\psi$ changes in permeabilized mouse embryonic fibroblasts (MEF) with intact Drp1 using 25 mM [$Na^+$] (**E**) Degree of depolarization (left) and the initial depolarization rate (right) for MEF. (**F**) Time-course of $Na^+$-uptake induced $\Delta\psi$ changes in permeabilized HeLa cells using 100 mM [$Na^+$]. (**G**) Degree of depolarization (left) and the depolarization rate (right) for HeLa cells. (**H**) Time-course of $Na^+$-uptake induced $\Delta\psi$ changes in MEF deficient in EMRE (EMRE-KO, left) or MCU/EMRE double KO (ME-KO, right) using 25 mM [$Na^+$]. (**I**) Summary data for degree of depolarization for EMRE KO (left), and MCU/EMRE double KO (ME-KO) (right) MEF. (**J**) Time-course of $Na^+$-uptake induced $\Delta\psi$ changes in HEK293 deficient in EMRE (EMRE-KO, left) or MCU/EMRE double KO (ME-KO, right) using 100 mM [$Na^+$]. (**K**) Summary data for degree of depolarization for EMRE-KO (left), and MCU/EMRE double KO (ME-KO, right) HEK293. **(L**) Time-course of $Na^+$-uptake induced $\Delta\psi$ changes in permeabilized hiPSCs using 100 mM [$Na^+$]. (**M**) Degree of depolarization (left) and the initial depolarization rate (right) for hiPSCs. Summary data presented as mean ± SEM; ****$P < 0.001$*, two-sample t-test, two-tailed, n=3-5.

### Figure S2.

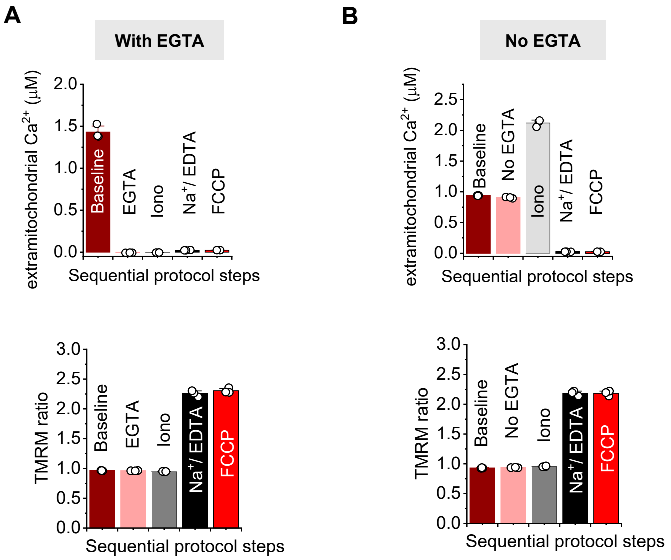


**Fig. S2. Matrix $Ca^{2+}$ depletion enhances MCUcx-mediated ion flux.**

(**A and B**) Extramitochondrial [$Ca^{2+}$] (Fura-8, Kd = 260 nM) measured at each step of the assay (upper panel) using isolated liver mitochondria in the presence of EGTA (A) or absence of EGTA (B) in the assay buffer. Note that Fura-8 was used here for reliable quantification of [$Ca^{2+}$] in the lower range but it saturates at ~2 μM as shown in panel B. Lower panel shows corresponding changes in Δψ (TMRM ratio) from the same experiment. Summary data presented as mean ± SEM, n=3.

**Figure S3.**

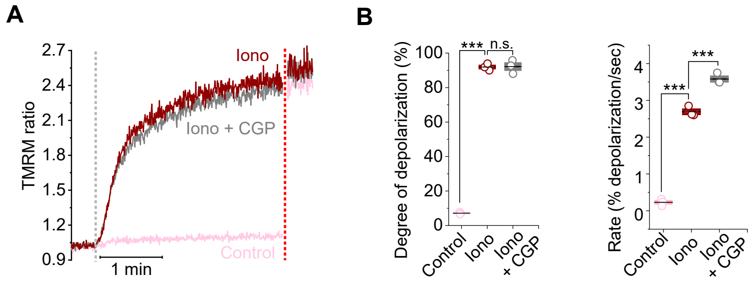


**Fig. S3. NDDR assay is not influenced by the inhibition of mitochondrial $Na^+/Ca^{2+}$ exchanger.**

(**A**) Time-course of $Na^+$-uptake induced Δψ changes in isolated liver mitochondria in the absence (Control) or presence of ionomycin (Iono), or in the presence of both ionomycin and CGP37157 (CGP), an inhibitor of mitochondrial $Na^+/Ca^{2+}$ exchanger (Iono + CGP). Grey dotted line denotes bolus injection of EDTA and $Na^+$; red dotted line marks FCCP addition. (**B**) Summary data for the degree of depolarization and the depolarization rate. Summary data presented as mean ± SEM; ****P < 0.001,* ns, not significant, One-way ANOVA with Bonferroni post hoc test, n=3.

**Figure S4.**

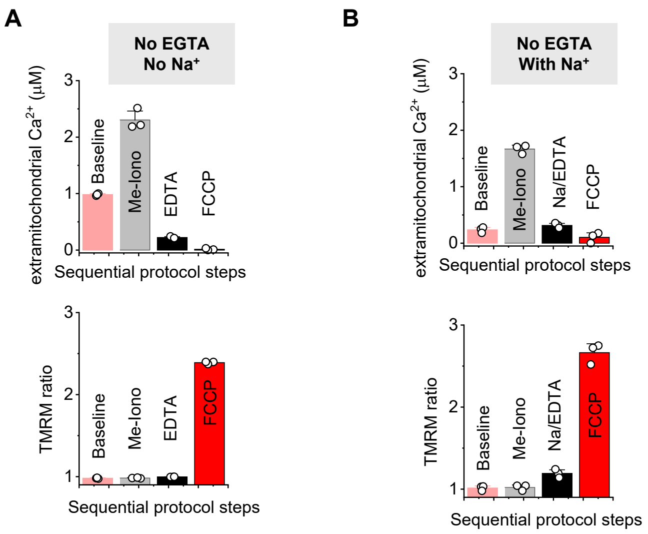


**Fig. S4. Non-chelating methyl ionomycin ester (Me-Iono) does not induce $Ca^{2+}$ release from mitochondrial subcompartments during the NDDR assay.**

(**A**) Extramitochondrial [$Ca^{2+}$] (using Fura-8FF) measured at each step of the assay (upper panel) when Me-Iono is used to replace Iono. Data are presented for direct comparison with the experiment shown in Fig. 2A. $Na^+$ is omitted from the assay. Upon Me-Iono application, minimal change in extramitochondrial [$Ca^{2+}$] (<2 μM) is observed implying inefficient release of $Ca^{2+}$ from mitochondrial subcompartments. Lower panel shows corresponding changes in $\Delta\psi$. (**B**) Extramitochondrial [$Ca^{2+}$] measured at each step of the assay (upper panel) in the presence of Me-Iono and $Na^+$. Data are presented for direct comparison with the experiment shown in Fig. 2A. $Na^+$ is unable to depolarize mitochondria when $Ca^{2+}$ is not depleted from the mitochondrial subcompartments Lower panel shows corresponding changes in $\Delta\psi$ (TMRM ratio). Isolated mouse liver mitochondria were used for these experiments. Summary data presented as mean ± SEM, n=3.

**Figure S5.**

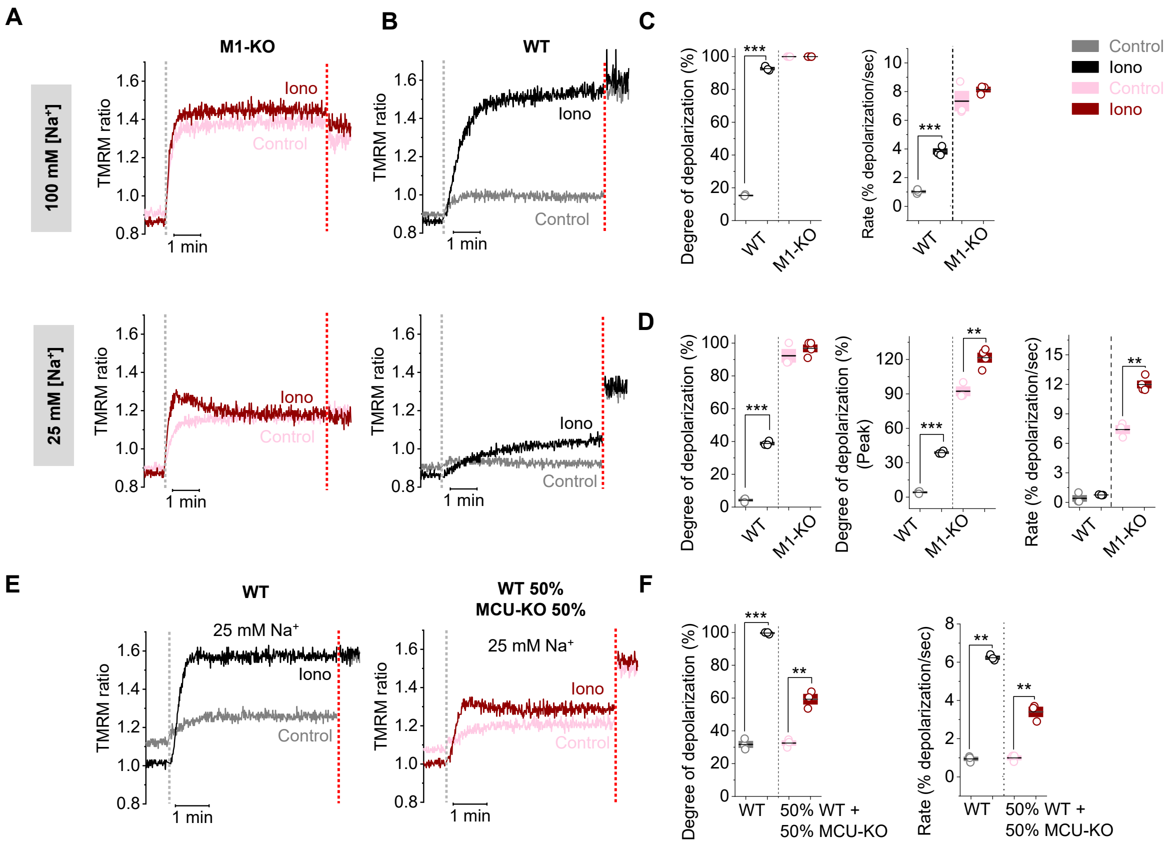


**Fig. S5. Matrix $Ca^{2+}$-dependent inhibition is functionally distinct from MICU1-mediated gatekeeping.**

(**A**) Time-course of $Na^+$-uptake induced Δψ changes in permeabilized MICU1-KO HEK293 (M1-KO) following addition of 100 mM (upper panel) or 25 mM [$Na^+$] (lower panel) in the absence (Control) or presence of ionomycin (Iono). (**B**) For comparison, representative time-courses for $Na^+$-uptake induced Δψ changes in WT cells are also shown under the respective conditions. (**C**) Degree of mitochondrial depolarization (left) and depolarization rate (right) of MICU1-KO and WT MEF in 100 mM [$Na^+$]. (**D**) Degree of mitochondrial depolarization at steady state (left), at peak (middle), and initial depolarization rate (right) of MICU1-KO and WT MEF in 25 mM [$Na^+$]. (**E**) Time-course of $Na^+$-uptake induced Δψ changes in permeabilized WT MEFs (WT, left), and a 1:1 mixture of WT and MCU-KO cells (WT 50% / MCU-KO 50%, right), following addition of 25 mM [$Na^+$] in the absence (Control) or presence of ionomycin (Iono). (**F**) Degree of mitochondrial depolarization (left) and depolarization rate (right) of WT MEFs (WT) and 1:1 mixture of WT and MCU-KO cells (WT 50% / MCU-KO 50%) in 25 mM [$Na^+$]. Note the ~50% reduction in the degree of depolarization in the 1:1 mixture of WT and MCU-KO cells, highlighting the need for stable and uniform expression across all cells for accurate interpretation. Grey dotted lines denote the bolus injection of EDTA and $Na^+$; red dotted lines mark FCCP addition. Summary data presented as mean ± SEM; *$P < 0.05$, **$P < 0.01$, ***$P < 0.001$, two-sample t-test, two-tailed, n=3-5.

### Figure S6.

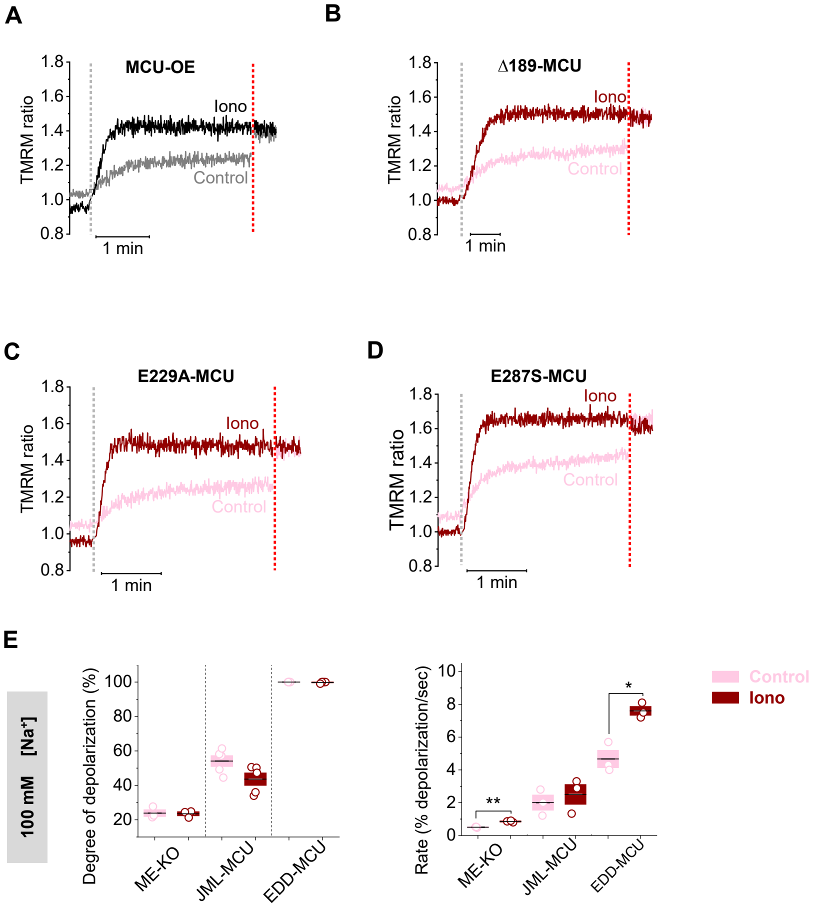


**Fig. S6. EMRE and a juxtamembrane loop in the MCU subunit are essential for MCI.** (**A-D**) Time-course of $Na^+$-uptake induced Δψ changes in MCU-KO MEFs stably expressing WT MCU (MCU-OE, trace is the same as depicted in fig. 5A for illustration, A), Δ57-189 MCU (Δ189-MCU, B), E229A-MCU (C) and E287S-MCU (D). Cells were exposed to 25 mM [$Na^+$] in the absence (Control) or presence of ionomycin (Iono). Grey dotted line denotes the bolus injection of EDTA and $Na^+$; red dotted line marks FCCP addition. (**E**) Degree of mitochondrial depolarization (left) and depolarization rate (right) for the respective EMRE-independent MCU variants stably expressed in MCU/EMRE double KO (ME-KO) MEFs. Cells were exposed to 100 mM [$Na^+$]. Summary data presented as mean ± SEM; ****P < 0.001,* two-sample t-test, two-tailed, n=3-5.
